# Computational and experimental characterization of the novel ECM glycoprotein SNED1 and prediction of its interactome

**DOI:** 10.1101/2020.07.27.223107

**Authors:** Sylvain D. Vallet, Martin N. Davis, Anna Barqué, Sylvie Ricard-Blum, Alexandra Naba

**Author notes:** **Corresponding authors:** Dr. Sylvie Ricard-Blum, Dr. Alexandra Naba.

## Abstract

The extracellular matrix (ECM) protein SNED1 has been shown to promote breast cancer metastasis and control neural crest cell-specific craniofacial development, but the cellular and molecular mechanisms by which it does so remain unknown. ECM proteins exert their functions by binding to cell surface receptors, sequestering growth factors, and interacting with other ECM proteins, actions that can be predicted using knowledge of protein’s sequence, structure and post-translational modifications. Here, we combined *in-silico* and *in-vitro* approaches to characterize the physico-chemical properties of SNED1 and infer its putative functions. To do so, we established a mammalian cell system to produce and purify SNED1 and its N-terminal fragment, which contains a NIDO domain. We have determined experimentally SNED1’s potential to be glycosylated, phosphorylated, and incorporated into insoluble ECM produced by cells. In addition, we used biophysical and computational methods to determine the secondary and tertiary structures of SNED1 and its N-terminal fragment. The tentative *ab-initio* model we built of SNED1 suggests that it is an elongated protein presumably able to bind multiple partners. Using computational predictions, we identified 114 proteins as putative SNED1 interactors. Pathway analysis of the newly-predicted SNED1 interactome further revealed that binding partners of SNED1 contribute to signaling through cell surface receptors, such as integrins, and participate in the regulation of ECM organization and developmental processes. Altogether, we provide a wealth of information on an understudied yet important ECM protein with the potential to decipher its functions in physiology and diseases.

## INTRODUCTION

The extracellular matrix (ECM) is a complex scaffold made of hundreds of proteins that instructs cell behaviors, organizes tissue architecture, and regulates organ function (1). It plays prominent roles during embryonic development, aging, and diseases (2–7). Mechanistically, ECM proteins can play these roles through their interactions with each other, with growth factors or morphogens, and with receptors present at the cell surface (1, 8, 9). These molecular interactions are mediated by specific protein domains, motifs, or sequences, and govern the nature of the chemical and mechanical signals conveyed by the ECM. Characterizing the composition of the ECM and the interactions taking place within this compartment, and determining how they regulate cellular functions is the first step towards building a systems biology view of the ECM.

Using the characteristic domain-based organization of known ECM proteins (10–12), we previously used sequence analysis to computationally predict the ensemble of genes coding for ECM proteins and ECM-associated proteins, and termed this ensemble the “matrisome” (13). Our interrogation of the human genome found that 1027 genes encoded matrisome proteins, among which 274 encoded “core” ECM components such as collagens, proteoglycans and glycoproteins. While all 44 collagen genes (14) and 35 proteoglycan genes (15) had previously been reported, several of the 195 genes predicted to encode structural ECM glycoproteins based on the protein domains present were or still are of unknown function (13, 16, 17). One such gene is *SNED1*. It encodes Sushi, nidogen and EGF-like domain-containing protein 1 (SNED1) and was originally named *Snep*, for stromal nidogen extracellular matrix protein, since the murine gene, *Sned1*, was cloned from stromal cells of the developing renal interstitium, and its pattern of expression overlapped with that of the ECM basement membrane proteins nidogens 1 and 2 (18). *Sned1* is broadly expressed during development, in particular in neural-crest-cell and mesoderm derivatives (18, 19), and interrogation of RNA-seq databases indicate that *SNED1* is also expressed in multiple adult tissues although at a very low level_1_. A decade after the cloning of this gene, we identified SNED1 in a proteomic screen comparing the ECM of poorly and highly metastatic mammary tumors and further reported the first function of this protein, as a promoter of mammary tumor metastasis (20).

Intrigued by this novel protein, we sought to identify its physiological roles. To do so, we generated a *Sned1* knockout mouse model and demonstrated that *Sned1* is an essential gene, since knocking it out resulted in early neonatal lethality (19). Furthermore, we showed that *Sned1* contributed to the formation of craniofacial and skeletal structures, since the few surviving knockout mice presented craniofacial abnormalities, were smaller in size, and had shorter long bones (19). However, the cellular and molecular mechanisms by which SNED1 regulates breast cancer metastasis or the development of craniofacial and skeletal structures remain unknown. Interrogation of the literature has revealed that the *SNED1* gene is found in large-omic datasets as dysregulated in different pathophysiological contexts or in response to various stimulations and in different *in-vivo* or *in-vitro* systems, hinting at a broader role for SNED1 (19). Yet, mining of gene expression, protein structure, and pathway databases revealed a paucity of data available on SNED1. This lack of tools and framework to study SNED1 creates an important gap in knowledge, given its pathophysiological significance.

Here, we used a combination of computational approaches and classical biochemical assays to analyze in depth the amino acid sequence and protein domains of SNED1, predict its secondary and tertiary structures and post-translational modifications, and build the first *ab-initio* 3D model of this protein and of its N-terminal region that contains a NIDO domain, a domain only found in 4 other human proteins. We further predicted its domain- and protein-level interaction networks which revealed the potential roles for SNED1 in multiple signaling pathways and biological processes. Altogether, our results offer the first molecular insights into this protein and a framework to start studying its likely multi-faceted roles in development, health, and disease.

## RESULTS

### Computational analysis of the sequence of the ECM protein SNED1

As previously described (18, 19), SNED1 is a multidomain protein containing one NIDO domain, one follistatin domain, one Sushi domain, also known as complement control protein (CCP) domain, 15 EGF-like and EGF-Ca_++_ domains, and 3 fibronectin III domains in its C-terminal region (**Figure 1A**, domain boundaries were predicted using SMART (21)). While these protein domains are found in lower organisms (22, 23), we have previously shown that SNED1 is only conserved among vertebrates (19).

**Figure 1.**
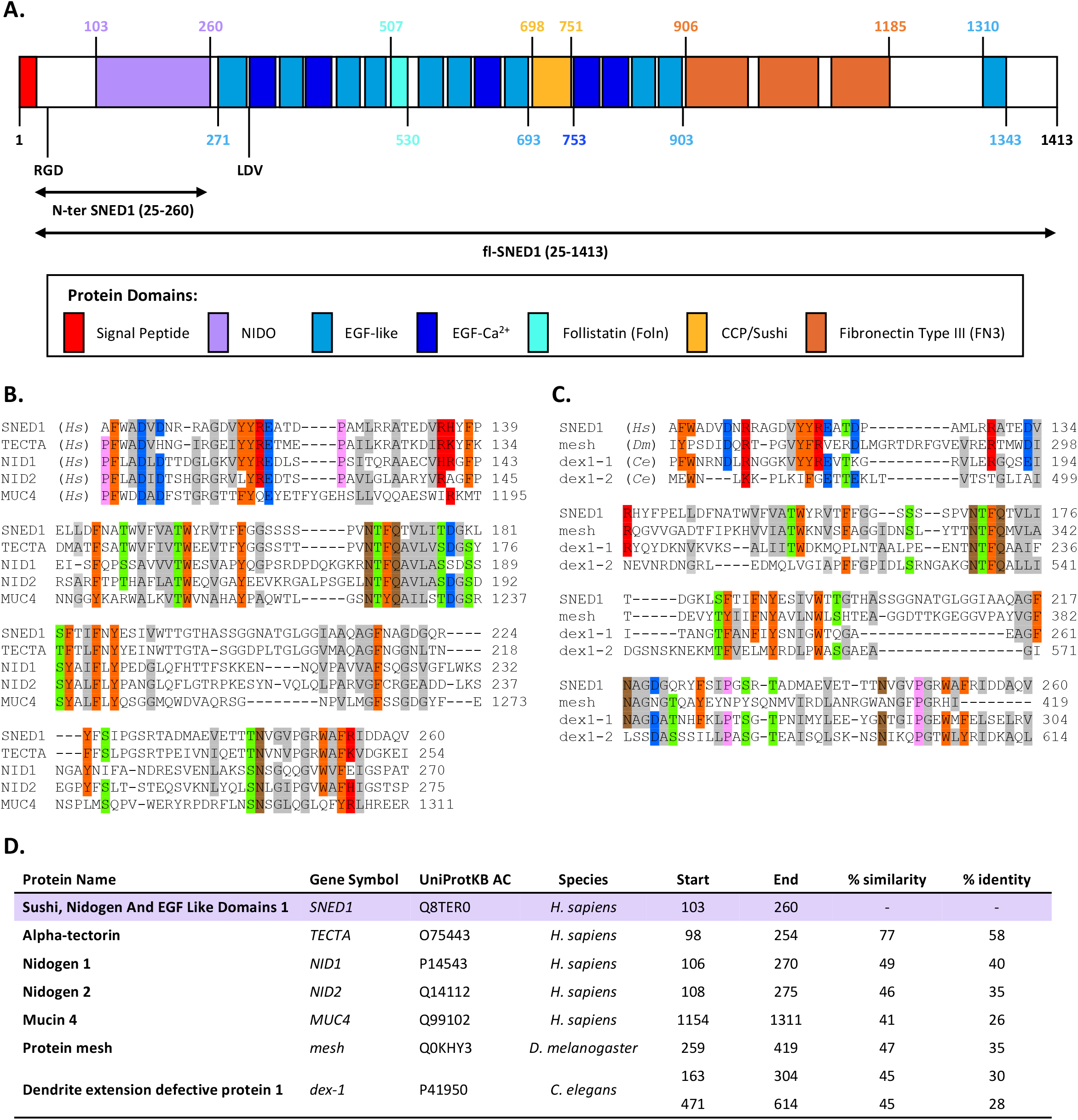
Domain-based organization of the ECM protein SNED1. **A.** Domain organization of human SNED1 (UniProtKB entry Q8TER0). Residues are numbered according to SMART (http://smart.embl-heidelberg.de/). Note that SMART prediction indicates that the second EGF-like domain resembles a follistatin domain. Potential integrin-binding sequences RGD and LDV are indicated. Arrows span the length of the recombinant proteins (N-terminal or fl-SNED1) used in this study. **B-D**. Sequence alignment of the NIDO domain of human (*Hs*) SNED1 with **(B)** the four human NIDO-domain-containing proteins: Alpha-tectorin (TECTA), Nidogen-1 (NID1), Nidogen-2 (NID2), and Mucin-4 (MUC4); and **(C)** two non-mammalian NIDO-domain-containing proteins: protein mesh (mesh, *D. melanogaster*), and dendrite extension defective protein 1 (dex1, *C. elegans*), containing 2 NIDO domains. **D**. Summary table in which the identity and similarity percentages are indicated for the NIDO domain of NIDO-containing proteins compared to the NIDO domain of SNED1.

An interesting feature of SNED1 is the presence of a NIDO domain (SMART: SM00539) in its N-terminal region (amino acids 103-260). This domain is only found in 4 other human or rodent proteins in addition to SNED1: the basement membrane proteins nidogen-1 and nidogen-2 (24, 25), mucin-4, and alpha-tectorin, a component of the tectorial membrane, the apical ECM of the inner ear (26–28). Identity between the NIDO domain of human SNED1 and that of other vertebrate SNED1 orthologues ranges between 73% and 92% (19). Sequence alignment within the NIDO domains of the 5 human proteins showed that the NIDO domain of SNED1 is most closely related to that of alpha-tectorin (77% of similarity and 58% of identity; **Figure 1B and 1D**). The NIDO domain is located in the N-terminal region of the 3 ECM proteins found in basement membranes (nidogens 1 and 2 and alpha-tectorin), while it is located in the center of mucin-4. The NIDO domain is also found in lower organisms used as model systems: the transmembrane protein mesh in *Drosophila* (29) and the secreted dendrite extension defective protein 1 in *C. elegans,* which contains two NIDO domains (**Figure 1C and 1D**). No function has clearly been associated with the NIDO domain, although the NIDO domain of mucin-4 was reported to regulate the ability of pancreatic tumor cells to invade and extravasate (30). In addition, we recently reported that the NIDO domain was one of the most mutated domains of ECM and ECM-associated genes in cancers (31). Understanding the molecular properties of this unique domain could thus provide insight into the function of SNED1 and its mechanism of action. Another remarkable feature of SNED1 is the presence of two possible integrin-binding motifs, RGD (amino acids 38-40) and LDV (amino acids 310-312), hinting at the fact that perhaps integrins are SNED1 receptors (*see below*).

### Development of a mammalian cell system to produce and purify SNED1 *in vitro*

In order to study the biochemical and biophysical properties of SNED1, we devised a mammalian cell system to produce recombinant FLAG-tagged full-length SNED1 (fl-SNED1) or the N-terminal fragment that contains the NIDO domain of SNED1 (**Figure 1A**). We found that both proteins were secreted by the cells, since we detected them in the conditioned medium of 293T cells stably expressing them, from which we can purify the proteins using the FLAG-tag added to their C-terminal ends (**Figure 2A**). In order to study SNED1, we also generated a rabbit polyclonal antibody, that we validated and found to be specific to human SNED1, since it did not recognize murine SNED1 (**Supplementary Figure S1A and S1B**).

**Figure 2.**
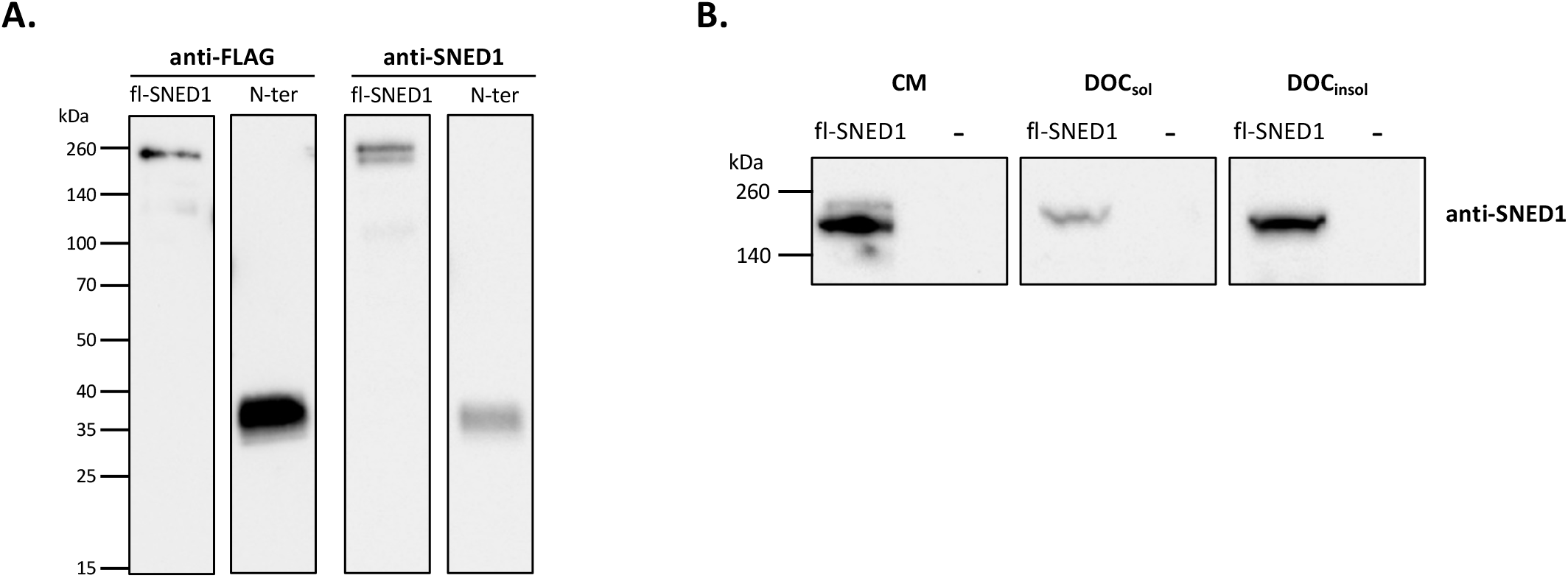
Expression of fl-SNED1 and of the N-terminal fragment in mammalian cells. **A.** Detection of fl-SNED1 and the N-terminal fragment of SNED1 by western blot using an anti-FLAG antibody and the anti-SNED1 polyclonal antibody described in this study (*see Supplementary Figure 1*). **B.** Deoxycholate (DOC) solubility assay on HEK293T cells expressing recombinant fl-SNED1 or an empty vector (-). Detection of fl-SNED1 in the conditioned medium (CM), deoxycholate-soluble (DOC_sol_) or DOC-insoluble (DOC_insol_) fractions by western blot using the anti-SNED1 polyclonal antibody.

The canonical deoxycholate (DOC) solubility assay has been used to demonstrate the incorporation of proteins, such as fibronectin, into the ECM (32, 33). Here, we show that fl-SNED is detected in the DOC-insoluble fraction, indicating its relative insolubility and likely incorporation in the ECM deposited by 293T cells *in vitro* (**Figure 2B**).

### Determination of the secondary and tertiary structures of full-length and of the N-terminal fragment of SNED1

Determining the structure of SNED1 has the potential to shed light on the mechanisms underlying its possible functions and signaling mechanisms. We thus turned to molecular modeling and biophysical assays using purified proteins to determine the secondary and tertiary structures of SNED1 and its N-terminal fragment.

#### Secondary structure of SNED1 and of its N-terminal fragment

The predicted secondary structure of the N-terminal fragment of SNED1 (amino acids 25-260) using Proteus2 (34) was 33% β-strand, 9.3% helix, and 57.6% random coil (**Supplementary Figure S2**), whereas the deconvolution of its circular dichroism (CD) spectra showed the presence of 39% β-strand, 5% helix, and 36% of random coil (**Figure 3A**). These results confirmed that the N-terminal fragmet of SNED1 containing the NIDO is mostly composed of β-strands, with a small percentage of helices (≤9%). Two helices were predicted in the NIDO domain itself (_124_PAMLRRATEDVRHY_137_ and _235_DMAEVETT_242_). A large proportion of random coil (73%) were predicted in fl-SNED1 together with 26% of β-strands, and 1% of helix corresponding to the sequence also found in the N-terminal fragment of SNED1, _124_PAMLRRATEDVR_135_ (**Figure 3A**). However, a higher percentage of β-strands (52%), and a lower amount of random coil (41%) together with 9% turns were found by CD analysis. Interestingly, no helix was experimentally detected in fl-SNED1.

**Figure 3.**
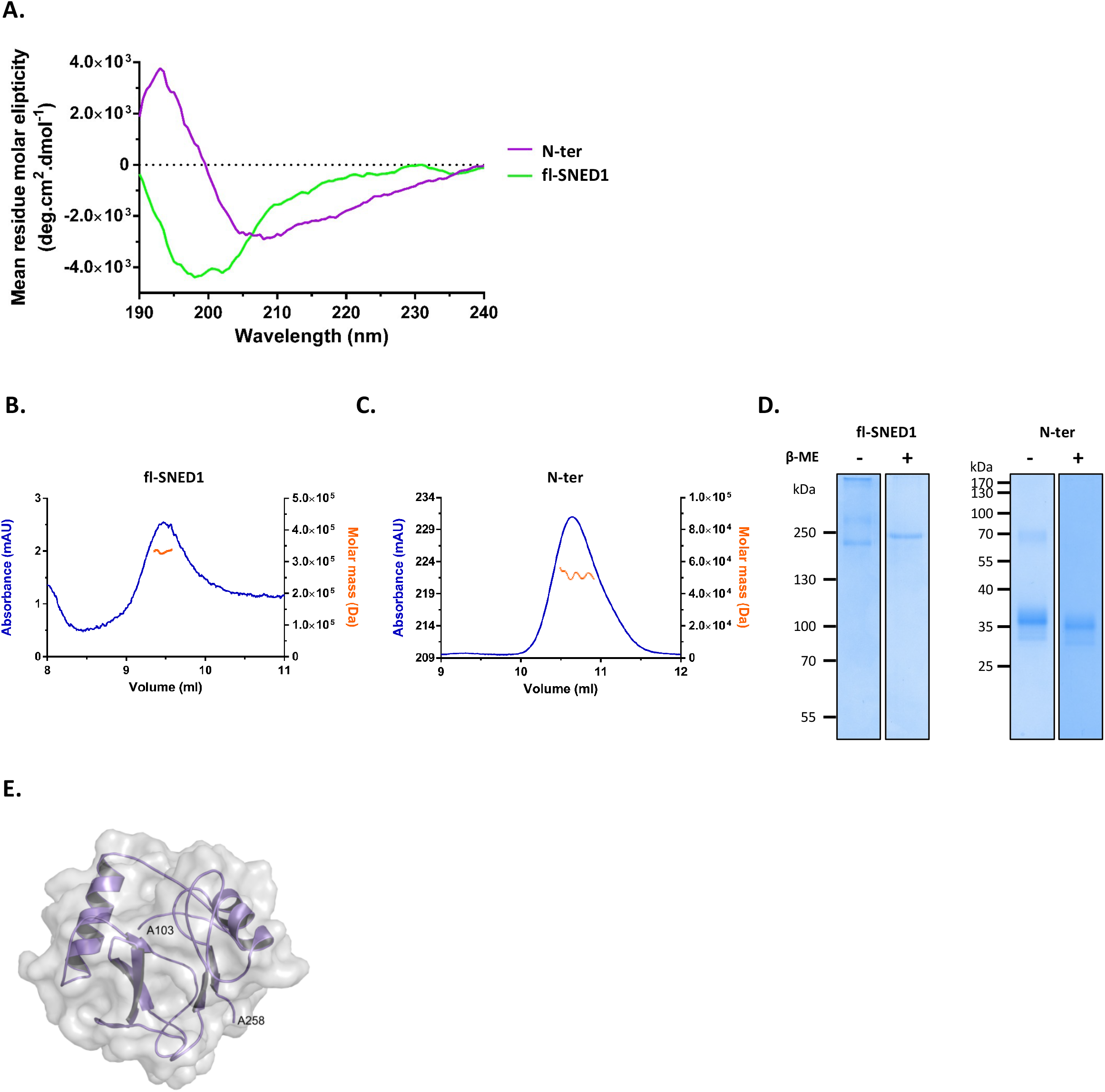
Determination of the molecular weight, secondary structure of fl-SNED1 and its N-terminal fragment, and *ab-initio* model of the NIDO domain. **A.** Circular dichroism spectra of human fl-SNED1 (green) and the N-terminal fragment of SNED1 (purple). **B-C.** Analysis of recombinant human fl-SNED1 (**B**), and N-terminal fragment of SNED1 (**C**) by Size-Exclusion Chromatography - Multi-Angle Laser Light Scattering (SEC-MALLS). **D.** Analysis of purified fl-SNED1 (left panel) and of its N-terminal fragment (right panel) by SDS-PAGE with and without reduction by β-mercaptoethanol (β-ME). **E**. *Ab-initio* model of the NIDO domain of human SNED1 generated by QUARK and refined with ModRefiner (TM-score 0.9, QMEAN −3.77).

#### Determination of the molecular weight of SNED1 and its N-terminal fragment using size-exclusion chromatography-multi-angle laser light scattering

The theoretical molecular weight (M_w_) of fl-SNED1 determined by the ProtParam tool (https://web.expasy.org/protparam/) is 150.8 kDa. Its absolute molar mass was determined to be 329 ± 5 kDa using size-exclusion chromatography-multi-angle laser light scattering (SEC-MALLS) (**Figure 3B**). The theoretical M_w_ of the N-terminal fragment of SNED1 is 26.6 kDa. Its absolute molar mass determined by SEC-MALLS was 51.7 ± 1.4 kDa (**Figure 3C**) and its M_w_ determined by size-exclusion chromatography (SEC) was 57 kDa (**Supplementary Figure S3**). These observations are consistent with the possible dimerization of SNED1 and its N-terminal fragment containing the NIDO domain, and with the presence of post-translational modifications (*see below*).

#### SNED1 and its N-terminal fragment are disulfide-bonded

The sequence of human SNED1 contains 107 cysteine residues. All the cysteine residues except one are located in two regions: residues 265-902 and 1311-1391. DISULFIND (35) predicted the presence of 53 disulfide bonds in the SNED1 sequence (**Supplementary Table S1**). Most domains of SNED1 were predicted to be disulfide-bonded except the fibronectin III domains (**Supplementary Table S1**). Only one cysteine residue, Cys_99_, is present in the N-terminus of SNED1.

To determine experimentally if SNED1 is stabilized by disulfide bonding *in vitro*, purified recombinant fl-SNED1 and its N-terminal fragment were analyzed by SDS-PAGE with or without the addition of the reducing agent β-mercaptoethanol (β-ME). Different migration profiles were observed on SDS-PAGE for fl-SNED1 depending on the presence or absence of reducing agent. Reduced SNED1 migrated as a single band with an apparent M_w_ of 210 kDa (*versus* 150.8 kDa for the theoretical M_w_), whereas an additional species (> 250 kDa) and higher Mw species, which did not enter the gel, were observed in the non-reduced condiction (**Figure 3D, left panel**). It is thus likely that unreduced SNED1 dimerizes *in vitro* and forms higher-Mw oligomers/aggregates stabilized by disulfide bonds.

In presence of reducing agent, the N-terminal fragment of SNED1 migrated with a higher apparent M_w_ (35 kDa) than the theoretical one (26.6 kDa) (**Figure 3D, right panel**). A molecular species migrating with an apparent M_w_ of 70 kDa was observed in absence of reducing agent, which suggests the existence of an intermolecular disulfide bond involving the only cysteine residue present within the N-terminal sequence.

#### Determination of the hydrodynamic radius of SNED1 and its N-terminal fragment

The hydrodynamic radius (Rh) of fl-SNED1 determined by dynamic light scattering (DLS) was 9.02 ± 0.97 nm. This value was more than two-fold higher than the value calculated for a fully folded protein comprising the same number of amino acid residues as SNED1 (3.7-3.9 nm), suggesting that SNED1 is an elongated protein, but was close to the Rh of an intrinsically disordered protein with the same residue number (10 nm), in agreement with the high percentage of random coil demonstrated by CD analysis (**Figure 3A**). We did not obtain a concentration of the N-terminal fragment of SNED1 sufficient enough to get a clear signal in the DLS experiment, and therefore a reliable experimental value of its hydrodynamic radius, but its theoretical hydrodynamic radius, calculated assuming that it folds as a globular protein, is 2.3-2.4 nm. The Stokes radius of the N-terminal fragment estimated by SEC was 3.4 nm. Altogether, our results provide the first computational and experimental determination of the structural and biophysical parameters of SNED1 (**Table 1**).

**Table 1.** Theoretical and estimated hydrodynamic parameters of SNED1.

| **Construct** | **Amino-acid Range** | **Sequence size**<br>**(incl. tag)** | **Theoretical $M_w$ ¶**<br>**(incl. tag)** | **Theoretical pI¶** | **Molar extinction coefficient**<br>**(M<sup>-1</sup>cm<sup>-1</sup>)¶** | **$R_g$**<br>**(nm)\*** | **$R_h$**<br>**(nm)\*** | **$R_h$**<br>**(nm)‡** | **DLS-derived $R_h$**<br>**(nm)** | **SEC-derived $R_s$**<br>**(nm)<sup>#</sup>** | **MALLS-derived $M_w$**<br>**(kDa)** |
| --- | --- | --- | --- | --- | --- | --- | --- | --- | --- | --- | --- |
| **Nter-FLAG** | 25-260 | 244 aa | 26.6 kDa | 4.7 | 40,910 | 1.8 | 2.3-2.4 | 2.4 | - | 3.4 | 52-55 |
| **Fl-SNED1-FLAG** | 25-1413 | 1397 aa | 150.8 kDa | 6.35 | S-S: 135,205<br>S-H: 128,580 | 3.4 | 3.7-3.9 | 4.4 | 9 | 4.5 | 329 |

¶ https://web.expasy.org/protparam/
\* Prediction based on sequence length using folded proteins parameters (62).
‡ Prediction based on theoretical $M_w$ (84).

### 3D model of the NIDO domain of SNED1 and full-length SNED1

Since no crystal structure or model of SNED1 or its NIDO domain are available, we sought to build a computational model. A model of the NIDO domain was built using QUARK and then refined with ModRefiner (coordinates file provided in **Supplementary File S1**). The TM-score was 0.9 (the topology is assumed to be correct if it is > 0.5), and the QMEAN, which should be above −4, was −3.77. It contained two helices, which were predicted by Proteus2 (although the second helix was longer than predicted, _233_TADMAEVETTT_243_), a short β-sheet and two additional β-strands (**Figure 3E**).

For fl-SNED1, we first employed two *ab-initio* protein structure prediction algorithms, I-TASSER (36) and QUARK (37), to generate 3D models exclusively based on SNED1’s amino acid sequence, but failed to build a reliable model of fl-SNED1. The best one was generated with I-TASSER and refined with ModRefiner (coordinates file provided in **Supplementary File S2**). The C-score, which should be comprised between −5 and 2, was −0.98, and the TM-score, which should be higher than 0.5, was 0.59; but the QMEAN was lower than −4 (−13.79) (**Supplementary Figure S4, upper panel**). Another model was generated with RaptorX (coordinates file provided in **Supplementary File S3**). The absolute global quality of this model was assessed by global distance test (GDT) and un-normalized GDT (uGDT), which should both be >50. The unrefined model had a uGDT of 597 but a GDT of 43 and a QMEAN of −4.66 (−4.87 for the refined model) (**Supplementary Figure S4, lower panel**). Both models predict that SNED1 is an elongated protein, which is consistent with the experimental Rh value (~9 nm) calculated by DLS (*see above*). While imperfect, these first models hint at an elongated structure in which multiple protein/protein interaction domains would be exposed and thus create docking sites for potential interactors.

### SNED1 is a glyco-phosphoprotein

A key feature of ECM proteins is their high level of post-translational modifications (PTMs), including glycosylations (38) and phosphorylations (39–41), with these modifications potentially further supporting protein-protein interactions and scaffolding (especially of mineralized tissues in the case of phosphorylations). We thus used several algorithms and queried multiple databases to determine whether SNED1 was predicted to be subject to PTMs (**Supplementary Table S2**) and further tested experimentally our findings.

#### Glycosylation and attachment of glycosaminoglycans

13 N-glycosylation sites were predicted in the sequence of human SNED1, including two sites in the N-terminal fragment of SNED1 at Asn_145_ and Asn_204_ (**Figure 4A** and **Supplementary Table S2A**) as well as 97 O-glycosylation sites (**Supplementary Table S2B**). dbPTM, the database compiling and curating experimentally verified PTMs, listed Asn_886_ and Asn_977_ as being N-glycosylated in murine SNED1 and Ser_1247_ as being O-glycosylated (**Supplementary Table S2D**). Of note, NetOGlyc did not predict the glycosylation of Ser_1247_ while it was detected experimentally. We thus sought to determine whether SNED1 produced in our mammalian culture system was glycosylated. Apparent M_w_ shift upon PNGase F treatment, which removes N-linked oligosaccharides, was calculated in three independent experiments. The treatment by PNGase F induced a downward shift of ~ 18 kDa in the apparent M_w_ of fl-SNED1 analyzed by western blot (**Figure 4B, left panel**), and of ~ 4 kDa in the apparent M_w_ of the N-terminal fragment of SNED1 (**Figure 4B, right panel**), demonstrating that both fl-SNED1 and its N-terminal fragment, when secreted by 293T cells, are N-glycosylated. We performed the same experiment using a mix of enzymes containing PNGase F and O-glycosidases but this did not result in a further downward shift of the SNED1 bands detected by western blot, suggesting that fl-SNED1 and its N-terminal fragment are not O-glycosylated when produced by 293T cells (*data not shown*).

**Figure 4.**
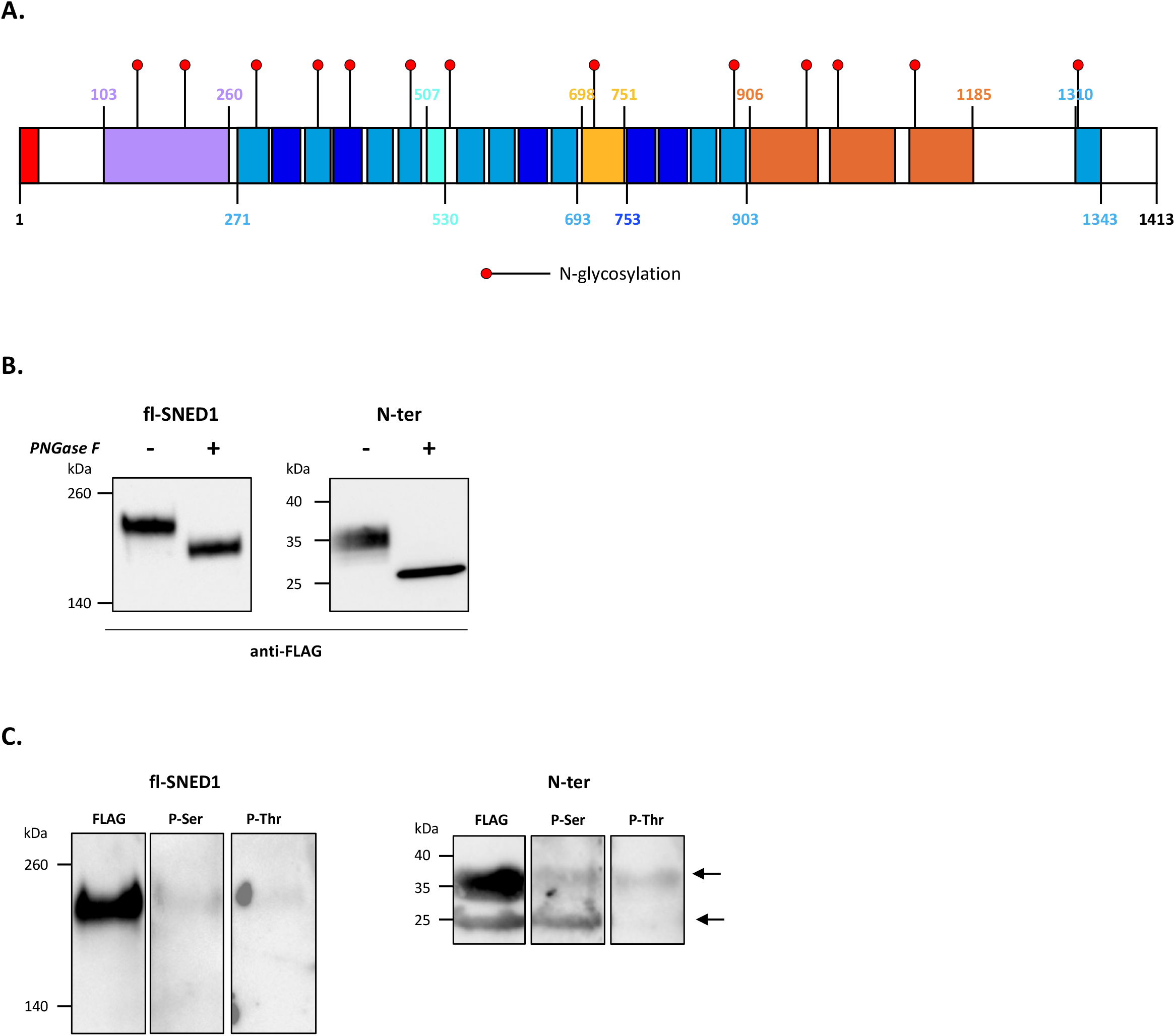
SNED1 is a glyco-phosphoprotein. **A.** SNED1 has 13 computationally predicted N-glycosylation sites (https://services.healthtech.dtu.dk/service.php?NetNGlyc-1.0) (*see complete prediction in Supplementary Table 3A*). **B.** Conditioned medium from cells expressing fl-SNED1 or the N-terminal fragment of SNED1 were treated or not with PNGase F and analyzed by western blot. **C.** Immunoprecipitation using the anti-FLAG affinity gel and analysis by western blot of immunoprecipitated fl-SNED1 and NIDO with anti-FLAG, anti-phosphoserine (P-Ser) or anti-phosphothreonine (P-Thr) antibodies.

In addition to glycosylation sites, several potential attachment sites of glycosaminoglycans (GAGs) were identified within the sequence of SNED1 including 8 Ser-Gly motifs, 1 Ser-Gly-X-Gly sequence (_846_SGGG_849_), and 2 Glu/Asp-X-Ser-Gly sequences, _66_EHSG_69_ and _918_EESG_921_ (**Supplementary Figure S5A**). However, the incubation of fl-SNED1 with the GAG-depolymerizing enzymes heparinase III or chondroitinase ABC did not induce a decrease in its M_w_ upon analysis by western blot, suggesting that no GAG chains are attached to recombinant human fl-SNED1 (**Supplementary Figure S5B**).

#### Phosphorylation

Sequence analysis also revealed 133 predicted phosphorylation sites in SNED1 (**Supplementary Table S2C**), including 22 residues within the N-terminal domain of SNED1, and 8 within the NIDO domain. However, it is worth noting that most of these sites are predicted by intracellular kinases not known to modify secreted ECM proteins (**Supplementary Table S2C**). However, interestingly, 6 residues are predicted to be phosphorylated by casein kinase II which is known to phosphorylate ECM proteins including collagen XVII (ecto-caseinase 2) fibronectin, and vitronectin (41).

Through database interrogation, we found experimental evidence showing the phosphorylation of twelve of these residues: 5 serine, 5 threonine, and 2 tyrosine residues, none of which lie within the N-terminal domain of SNED1 (**Supplementary Table S2D**). In order to determine whether SNED1 was phosphorylated when secreted by 293T cells, we immunoprecipitated FLAG-tagged SNED1 and conducted western blot analysis of the immobilized protein using anti-phosphoserine, anti-phosphothreonine, and anti-phosphotyrosine antibodies. While we were not able to obtain consistent results with the anti-phosphotyrosine antibody, our results show that both human fl-SNED1 (**Figure 4C, left panel**) and its N-terminal fragment (**Figure 4C, right panel**) were phosphorylated on serine and threonine residues. Of note, we observed two bands when we immunoprecipitated the N-terminal fragment of SNED1 with the anti-FLAG antibody and, while both were detected with the anti-phosphothreonine, only the lower band showed a signal with the anti-phosphoserine.

Altogether our results provide evidence that SNED1 secreted by 293T cells is both N-glycosylated and serine- and threonine-phosphorylated, and that some of the modified residues lie within the N-terminal region of SNED1. Future studies are needed to determine which enzymes are responsible for these PTMs and how these PTMs relate to SNED1 structure, interactions and functions.

### Domain-domain interaction network of SNED1

A domain-domain interaction network of fl-SNED1 was built using 3did, the database of 3-dimensional interacting domains, and returned 106 unique interactions (**Figure 5** and **Supplementary Table S3**) when queried for domains present in SNED1. 49 domains were predicted to bind to the fibronectin type III domains, 24 to the sushi domain, 22 to EGF-like domains, 11 to calcium-binding EGF-like domains, and 2 to the follistatin domain (**Figure 5**). Two interactions were retrieved twice (Sushi/EGF and EGF/EGF_CA). Of note, the lack of knowledge about the NIDO domain was further exemplified here, since NIDO is absent from the database 3did and no other protein domain has ever been reported to interact with it.

**Figure 5.**
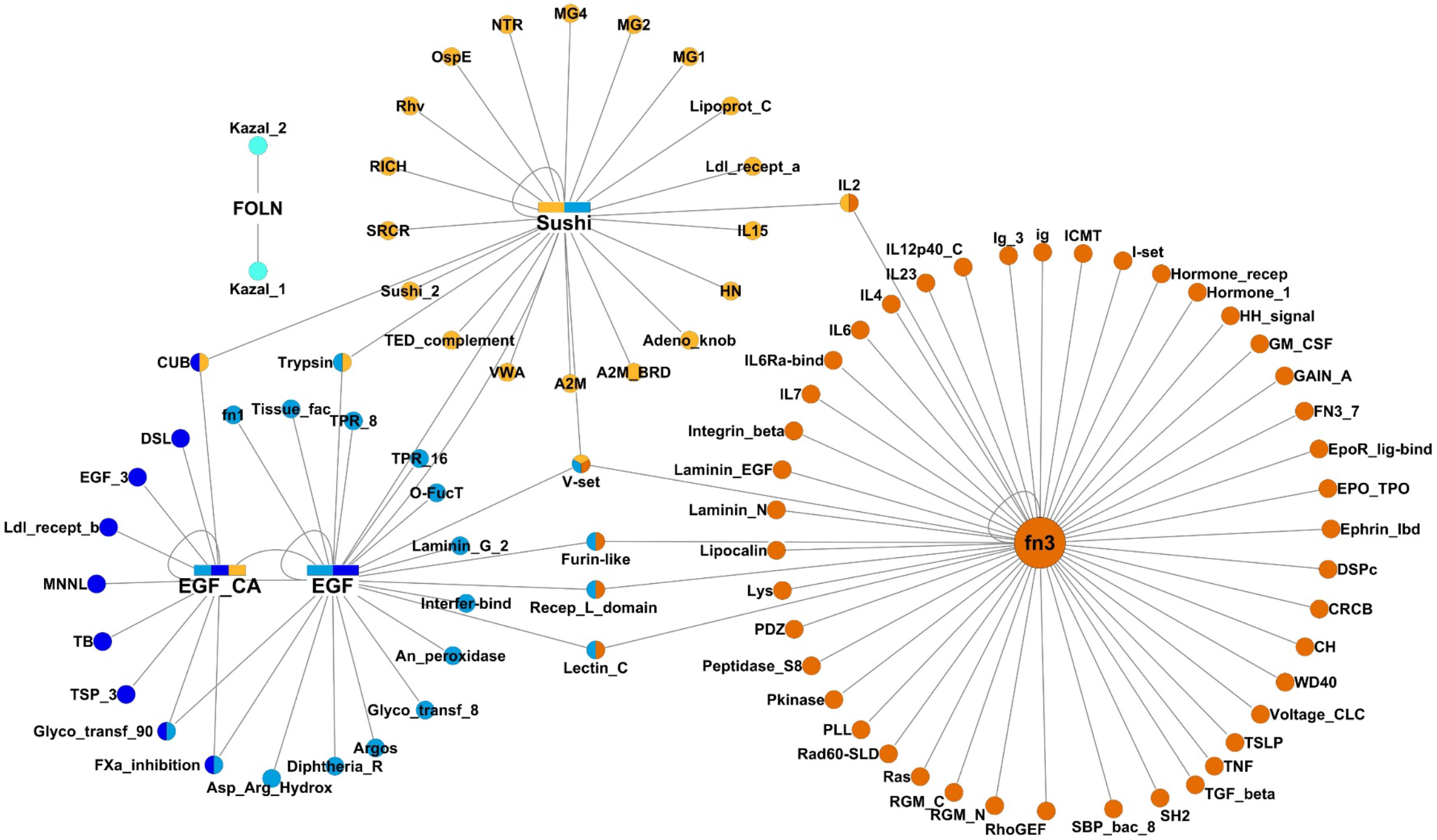
Domain-domain interaction network of SNED1. Predicted domain-domain interaction network of SNED1. Domains are colored according to the domains of SNED1 they interact with (FOLN: cyan, EGF: blue, EGF_CA: dark blue, fn3: orange, Sushi: light orange).

### Prediction of the protein-level SNED1 interactome

The query of the interaction databases MatrixDB (42) and IntAct (43) returned only one partner of SNED1, the estrogen receptor beta (44) (*see below*). This prompted us to use computational approaches to predict interactions established by SNED1 with other proteins (*see Materials and Methods*). We focused on secreted proteins and membrane proteins, which resulted in the prediction of 114 unique interactions by at least one algorithm, including SNED1 auto-interaction (**Figure 6A**). More than half of the protein partners of SNED1 were annotated as membrane proteins in UniProtKB (66/114, 57.9%). Of these, 11 (ITGA1, ITGA10, ITGA11, ITGA4, ITGA7, ITGB1, ITGB2, ITGB3, ITGB4, ITGB5, ITGB7) were integrins, which constitute the major class of ECM receptors (45). Sequence analysis of SNED1 revealed that it displays two integrin-binding consensus sequences, RGD and LDV (**Figure 1A**), which suggests that integrins may serve as SNED1 receptors. Two additional membrane proteins, Indian hedgehog (IHH) and tissue factor (F3), are also annotated as matrisome-associated and secreted proteins respectively (**Figure 6A** and **Supplementary Table S4**). Last, 47 partners of SNED1 are identified as being extracellular proteins, including 30 matrisome proteins, 10 matrisome-associated proteins, and 7 secreted proteins (**Figure 6A** and **Supplementary Table S4**). Among the 30 matrisome proteins are the 6 collagens: COL6A3, found in basement membranes and other ECMs, COL7A1, and the Fibril-Associated Collagens with Interrupted triple-helices (FACITS (14)), all containing a thrombospondin domain, COL12A1, COL14A1, COL16A1, COL20A1); and a number of ECM glycoproteins: 4 tenascins (TNC, TNN, TNR, and TNXB) (46), fibronectin (FN1), the latent-TGFβ binding protein 2 (LTBP2), and the basement membrane glycoproteins nidogens 1 and 2. 45 unique interactions were also predicted to involve intracellular proteins (**Supplementary Table S4**).

**Figure 6.**
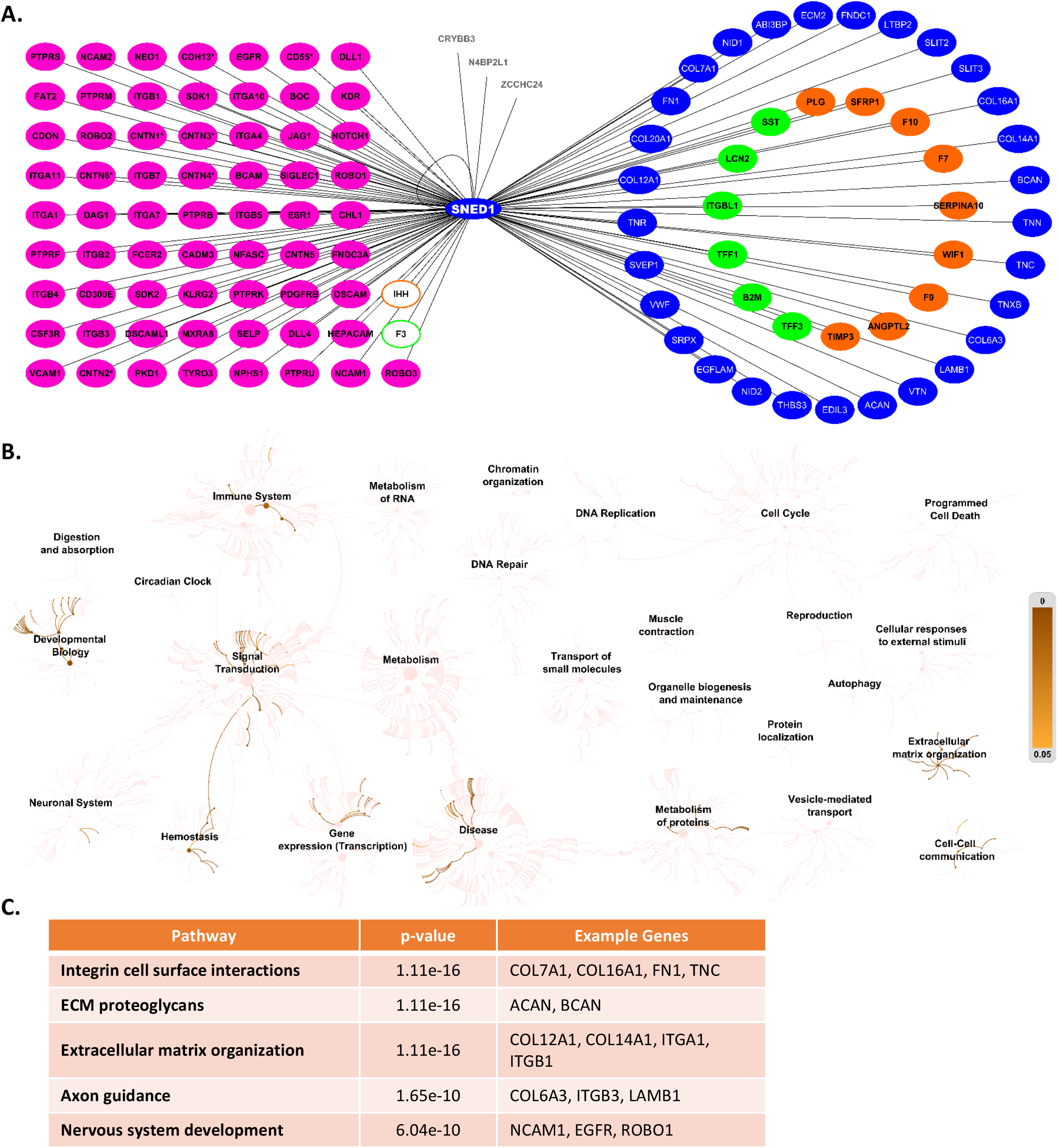
Prediction of the SNED1 interactome and pathway analysis. **A.** Predicted extracellular and membrane protein-protein interaction network (interactome) of human SNED1. Proteins are color-coded according to their location (blue: core matrisome, orange: matrisome-associated, green: secreted, pink: membrane). Proteins with undetermined locations are in grey. (*) Glycosylphosphatidylinositol-anchored protein. **B.** Genome-wide overview of the processes potential SNED1 interactors contribute to built with Reactome. Darker color indicates more representation of interaction within that pathway. Each step away from the center of a pathway represents one level lower in the pathway hierarchy. **C.** Top five cellular pathways in which the SNED1 interactome was found to be significantly enriched, sorted by p-value.

Comparison of the predictions made by the different algorithms revealed that 10 interactions were predicted by at least two methods including the ECM proteins collagen VII (COL7A1), tenascin N (TNN), and fibronectin (FN1), the ECM receptor integrin β4 subunit (ITGB4) and the related secreted integrin β-like protein-1 (ITGBL1), the transmembrane proteins basal cell adhesion molecule (BCAM), NOTCH1, the cell adhesion molecule-related/down-regulated by oncogenes (CDON), polycystin-1 (PKD1), and the signaling molecule Indian hedgehog protein (IHH) (**Supplementary Figure S6A** and **Supplementary Table S4**).

We then focused on the 13 binding partners predicted to interact with SNED1 by HOMCOS (**Supplementary Table S4E**). This tool specifically allows the 3D modeling structures and interactions using 3D molecular similarities, resulting in the mapping of the potential interactor binding sites to SNED1 domains. We found that most putative interactor binding sites were located within EGF-like domains, whereas only a few partners, including the proteoglycan aggrecan (ACAN) and the ECM glycoprotein fibronectin (FN1), were predicted to interact with the fibronectin III domains (**Supplementary Figure S6B**). Interestingly, three glycosylation enzymes, the protein O-glucosyltransferase 1 (POGLUT1), alpha-1,3-xylosyltransferase (XXYLT1) and protein O-fucosyltransferase 1 (POFUT1), were predicted to bind SNED1 (**Supplementary Figure S6C**). These enzymes have previously been shown to glycosylate residues in EGF-like repeats of transmembrane proteins, in particular in Notch or Notch ligands (e.g. JAG1), or of secreted proteins (47–50). While our experiments have shown that SNED1 is not O-glycosylated in our cellular system, the Ser_1247_ has been shown to be modified by N-Acetyl-D-galactosamine in MCF-7 and MDA-MB-231 cells (51) and the above enzymes are thus potential candidates to catalyze SNED1 O-glycosylation.

Of note, no partner was predicted to interact with the NIDO, follistatin or C-terminal domains of SNED1 (**Supplementary Figure S6B and C**), which, again, may reflect the limited experimental data available for these domains.

### Potential binding partners of SNED1 are involved in multiple signaling pathways

The *in-silico* interaction network of SNED1 was then analyzed using Reactome (52) to identify associated biological pathways. No annotation could be retrieved for SNED1 itself, further highlighting the critical gap in knowledge about this protein (*Reactome, version 73, released June 17, 2020*). The Reactome database included information on 94 of the 114 predicted SNED1 partners, and since the majority of them are either part of the matrisome or are transmembrane receptors, the processes most over-represented in SNED1’s network were “signal transduction”, “cell-cell communication”, “ECM organization” and “developmental biology” (**Figure 6B**). More specifically, the predicted SNED1 interactors were found to be part of, or to contribute significantly to, more defined pathways including “integrin cell surface interaction” and “ECM proteoglycans” (**Figure 6C**). This analysis, together with the list of predicted SNED1 interactors, will help prioritize future lines of investigation focused on uncovering the molecular mechanisms by which SNED1 controls aspects of embryonic development (19) and breast cancer metastasis (20).

## CONCLUSION

Deciphering the nature of protein-protein interactions within the ECM is critical to understand the mechanisms governing proper ECM assembly and signaling functions in health and disease. We report here the computational prediction of the structure and interaction network of the novel ECM protein SNED1 and provide experimental insight on the properties of this protein. While SNED1 shares structural and biochemical features with other known ECM components, its mechanistic roles remain unknown. And while our study revealed certain characteristics of this protein, it has also raise multiple novel questions, for example, is the dimerization of SNED1 physiological, and if so, what function does it play? What are the functional roles of SNED1 glycosylation and phosphorylation?

The NIDO domain found in the N-terminal region of SNED1 is only found in 4 other human proteins. Structure/function analysis of the NIDO domain of mucin-4 has revealed its role in promoting invasiveness of pancreatic tumor cells (30, 53). We have previously demonstrated that SNED1 promotes mammary tumor metastasis (20), and SNED1 was also identified in a screen as a mediator of p53-dependent pancreatic tumor cell invasive phenotype (54). It would thus be interesting to conduct a detailed structure/function study and determine whether SNED1 promotes tumor cell invasiveness via its NIDO domain, and if so, what are the specific mechanisms at play. In addition, since 3 of the 4 NIDO-domain-containing proteins (nidogens 1 and 2 and alpha-tectorin) are basement membrane proteins, it would be interesting to employ electron microscopy on tissues expressing SNED1 to determine whether it also localizes to basement membranes.

Our study has also identified over one hundred putative binding partners of SNED1. Among these, are 11 integrin chains or subunits (5 α chains and 6 β chains), a major class of ECM receptors and key players in cell functions in health and diseases, including cancers (45, 55). In humans, 18 α subunits and 8 β subunits can assemble to form 24 different heterodimers (45), among which, 8 can recognize the RGD motif (α5β1, α8β1, αvβ1, αvβ3, αvβ5, αvβ6, αvβ8, and the platelet-specific αIIbβ3) and 3 can recognize the LDV motif (α4β1, α9β1, and α4β7, the latter being found at the surface of hematopoietic cells). With SNED1 presenting both an RGD motif and an LDV motif, it would be interesting to determine experimentally if integrin heterodimers can indeed bind SNED1 and if so, which signaling pathways and cellular functions are triggered downstream of the SNED1/integrin interaction.

We have previously observed that knocking down *SNED1* in mammary tumor cells resulted in a significant decreased in metastasis and that *SNED1*-knockdown mammary tumors were surrounded by a thick capsule of fibrillar collagens (20). Analysis of the functions to which putative SNED1 interactors contribute identified “ECM organization” as one of the most significantly enriched pathway, and 6 collagen chains (COL6A3, COL7A1, COL12A1, COL14A1, COL16A1, and COL20A1) were predicted to bind to SNED1. Together, these results hint at a potential role for SNED1 in regulating collagen deposition and organization, which will need to be experimentally assessed.

While our focus is on the full-length, secreted ECM protein SNED1, early reports indicated that the 3’ half of *SNED1* could encode an intracellular protein then named insulin response element-binding protein 1 (IRE-BP1), since it was identified by phage display to bind the IRE of the gene encoding insulin-like growth factor-binding protein 3 (IGFBP3) (56, 57). Sequence analysis suggests that an alternative start codon could generate this shorter isoform, although no experimental evidence supports this. Interestingly, while none of the predicted interactors found via molecular modeling in our study have been validated experimentally, SNED1 was found to interact, by mass spectrometry, with the intracellular estrogen receptor beta (44). Whether the full-length secreted SNED1 can interact with this protein, or whether it is an intracellular shorter isoform or a truncated form of SNED1 remains to be determined, as does the physiological relevance of this interaction.

In summary, our study has provided the first biochemical and biophysical insights into the novel ECM protein SNED1 and is paving the way for future mechanistic studies that will eventually help us understand its multi-faceted roles in development, health, and disease.

## EXPERIMENTAL PROCEDURES

### Plasmid constructs

The cDNA encoding full-length human SNED1 (fl-SNED1) cloned into pCMV-XL5 (clone SC315884) was obtained from Origene (Rockville, MD). The cDNA encoding full-length murine *Sned1* cloned into pCRL-XL-TOPO (clone 40131189) was obtained from Open Biosystems (now, Thermo Fisher, Waltham, MA). Fl-SNED1 and a construct spanning the most N-terminal region of SNED1 and including the NIDO domain (amino acids 1 to 260, referred to as “N-terminal fragment of SNED1” in the text and as “N-ter” in the figures) were subcloned into the bicistronic retroviral vector pMSCV-IRES-Hygromycin between the *Bgl*II and *Hpa*I sites, and a FLAG tag (DYKDDDDK) was added at the C-terminus of the constructs (**Figure 1A**). These constructs were used to establish stable cell lines (*see below*). 6x-His-tagged constructs of human and murine SNED1 cloned into pCDNA5/FRT (Thermo Fisher, Waltham, MA) between the *FseI* and *AscI* sites were used to transiently transfect 293T cells to validate the anti-SNED1 antibody generated in this study (**Supplementary Figure S1**). All primers used are listed in **Supplementary Table S5**. All constructs were validated by Sanger sequencing.

### Expression of full-length SNED1 and the N-terminal fragment of SNED1 in mammalian cells and purification strategy

Human embryonic kidney (HEK) 293T cells (further referred to as 293T cells) were cultured in Dulbecco’s Modified Eagle’s medium (DMEM) supplemented with 10% fetal bovine serum and 2 mM glutamine.

#### For retrovirus production

Cells were plated at ~30% confluency and transfected 24 h later using the Lipofectamine 3000 system (Invitrogen) with a mixture containing 1 μg of retroviral vector with the construct of interest, 0.5 μg of packaging vector (pCL-Gag/Pol), and 0.5 μg of coat protein (VSVG). The transfection mix was prepared using the manufacturer’s instructions and added to the cells for 24 h, after which the transfection mix was removed and cells were fed with fresh culture medium. Cells were then cultured for an additional 24 h, after which the viral-particle-containing culture medium was collected and filtered through a 0.45-μm filter and then either stored at −80 °C or immediately used.

#### To establish cells stably expressing fl-SNED1 or the N-terminal fragment of SNED1

Cells were plated at ~40% confluency. Undiluted viral-particle-containing conditioned medium was added to the cells 24 h after seeding, and cells were fed with fresh culture medium 24 h after transduction. Transduced cells were selected with hygromycin (100 μg/ml) over a period of 10 days.

Protein expression and secretion were monitored by performing western blot analysis on cellular protein extracts obtained by lysing cells using 3X Laemmli buffer (0.1875 M Tris-HCl, 6% SDS, 30% Glycerol) supplemented with 100 mM dithiotreitol, and on the cell conditioned media (CM) with the rabbit polyclonal anti-SNED1 antibody (2 μg/mL) described below, a rabbit polyclonal anti-FLAG antibody (2 μg/mL; Sigma, F7425), or the monoclonal anti-FLAG M2 antibody (Sigma, F3165). Secondary anti-rabbit antibody conjugated to the horseradish peroxidase (Thermo Fisher, 31460) was used and immunoreactive bands were detected by chemiluminescence (SuperSignal West Pico PLUS, Thermo Fisher or ECL Prime Western Blotting System, GE Healthcare).

For large-scale expression of fl-SNED1 and the N-terminal fragment of SNED1, cells were cultured in HYPER*Flasks*™ in DMEM (Sigma-Aldrich, D5796) supplemented with 50 μg/ml of gentamicin (Sigma-Aldrich, G1272) as previously described (58). Culture media were harvested every 48 h for up to 18 days. After collection, 3 tablets of EDTA-free cOmplete inhibitor (Roche) were added to the media which was then centrifuged at 14,000 × *g* for 30 min at 4°C. Supernatants were stored at −80°C until use. Fl-SNED1 and the N-terminal fragment were purified by affinity chromatography on an anti-FLAG resin (Sigma-Aldrich, A2220) as previously described (58) in presence of 150 mM NaCl. In brief, FLAG-tagged proteins were purified at 4°C on the anti-FLAG resin at a flow rate of 20 ml/h with a P1 pump (GE Healthcare), and eluted by competition with a FLAG peptide solution at 200 μg/ml in 10 mM Hepes, 150 mM NaCl, pH 7.4 (HBS). The purified proteins were then concentrated on either Amicon (Merck Millipore, MWCO 10 kDa) or Vivaspin (Sartorius, MWCO 5 kDa) concentration columns.

### Generation of a rabbit polyclonal anti-SNED1 antibody

A polyclonal antibody directed against human SNED1 was generated against a SNED1 peptide spanning amino acids 29 to 41 (_29_ADFYPFGAERGDA_41_, **Supplementary Figure S1**) with a cysteine residue at the C-terminus. Peptide synthesis, conjugation, rabbit immunization and serum collection were performed by Covance (Denver, PA, USA) and the initial step of development performed in the laboratory of Dr. Richard Hynes (MIT). The antibody was purified by affinity chromatography using the peptide immobilized on an agarose resin via the Sulfo-Link chemistry according to the manufacturer’s instruction (Thermo Scientific). The antibodies were eluted sequentially with a 0.2 M glycine solution at pH 3 and pH 2.5, dialyzed against phosphate-buffered saline (PBS) and stored at 4°C. The reactivity and specificity of the antibody were assessed by western blot (**Supplementary Figure S1**).

### DOC solubility assay

Cells stably expressing FLAG-tagged fl-SNED1 were grown to confluency and lysed in DOC buffer: 2% deoxycholate; 20 mM Tris-HCl, pH 8.8 containing 2mM EDTA, 2mM N-ethylamine, 2mM iodoacetic acid, 167 μg/mL DNase and 1X protease inhibitor (Thermo Scientific, A32953) as previously described (32). Lysate was then passed through a 26G needle to further shear DNA and reduce viscosity. Centrifugation was used to pellet the DOC-insoluble, ECM-enriched, protein fraction from the DOC-soluble supernatant, enriched for intracellular components. These fractions were analyzed for the presence of SNED1 by western blot as described above.

### Analysis of SNED1 post-translational modifications by SDS-PAGE and western blot

Conditioned media from 293T cells stably expressing FLAG-tagged fl-SNED1 and the N-terminal fragment of SNED1 were incubated with PNGase F as previously described (58), or with heparinase III and chondroitinase ABC (2 mU per 40 μL of CM) as previously described (59). Proteins were separated by SDS-PAGE, and transferred onto nitrocellulose membranes. Membranes were probed with an anti-FLAG antibody or the rabbit polyclonal anti-SNED1 antibody generated in this study to identify recombinant SNED1.

To determine whether SNED1 is phosphorylated, FLAG-tagged fl-SNED1 was immunoprecipitated from the conditioned medium of cells stably expressing this construct on the anti-FLAG resin (Sigma-Aldrich, A2220), bound proteins were resolved by SDS-PAGE and western blots were performed with the anti-FLAG antibody to validate the immunoprecipitation of FLAG-tagged fl-SNED1 and with anti-phosphoserine (1 μg/mL; Abcam, ab9332), anti-phosphothreonine (1 μg/mL; Sigma-Aldrich, AB1607), or anti-phosphotyrosine (1 μg/mL; Sigma-Aldrich, 05-321) antibodies.

### Circular dichroism (CD)

Far-UV CD spectra were recorded in a quartz cuvette at 20°C with a path length of 0.1 cm in a Chirascan spectrometer (Applied Photophysics, UMS3444/US8, Protein Science Facility, Lyon, France), from 260 to 180 nm with a step of 0.5 nm, a response time of 1 s and a bandwidth of 0.5 nm for the N-terminal fragment of SNED1 and 1 nm for fl-SNED1. Fl-SNED1 and the N-terminal fragment of SNED1 were dialyzed *versus* 10 mM potassium phosphate buffer pH 7.4 using Micro DispoDialyzers (MWCO 10 kDa). Five spectra were collected for the N-terminal fragment of SNED1 (6.6 μM) and SNED1 (0.89 μM) and the buffer signal was subtracted. The spectra were deconvolved using DichroWeb (60), the CDSSTR program (61) and the reference data sets 7 and SP175 for the N-terminal fragment of SNED1 and fl-SNED1 respectively.

### Size-Exclusion Chromatography - Multi-Angle Laser Light Scattering (SEC-MALLS)

Human fl-SNED1 and the N-terminal fragment of SNED1 were analyzed by SEC-MALLS on the SWING beamline at SOLEIL synchrotron (Saint-Aubin, France) at 20°C with a Wyatt Dawn Heleos light scattering instrument equipped with an Optilab rEX refractometer and a spectrophotometer for molecular mass measurements. 50 μl of the N-terminal fragment of SNED1 (13.2 μM) and fl-SNED1 (4.5 μM) were injected on Superdex 75 Increase 10-300 GL and Superdex 200 Increase 10/300 GL columns (GE Healthcare) respectively at a flow rate of 0.5 ml/min at 20°C, see Figure 3B and 3C. Data were analyzed with ASTRA software 6.1.7 (Wyatt Technology).

### Dynamic Light Scattering (DLS)

Three series of ten acquisitions were performed at 20°C and an angle of 90° for fl-SNED1 at a concentration of 4.5 μM in HBS on a Zetasizer μV (Malvern Instruments Ltd, SOLEIL synchrotron, Saint-Aubin, France) in a disposable plastic cuvette. Data were analyzed with the Zetasizer software 7.12 (Malvern Instruments Ltd). The theoretical hydrodynamic radii of fl-SNED1 and the N-terminal fragment of SNED1 were calculated using folded proteins parameters and the number of amino acid residues of each protein (62).

### Bioinformatic analysis of the amino acid sequence of human SNED1

The sequence of human SNED1, without its peptide signal, was used for further queries unless stated otherwise (UniProtKB Q8TER0, residues 25-1413; **Figure 1A**).

The domain organization of human SNED1 was drawn with Illustrator of Biological Sequences 1.0.3 (63). The secondary structure of SNED1 and NIDO was predicted using Proteus2 (34). Multiple sequence alignment of the NIDO domains of SNED1 and other mammalian and non-mammalian proteins was performed using Clustal Omega (https://www.ebi.ac.uk/Tools/msa/clustalo/) and identical or similar residues in over 70% of the sequences were color-coded with Color Align (https://www.bioinformatics.org/sms2/color_align_prop.html). Pairwise comparison was performed with Protein BLAST (https://blast.ncbi.nlm.nih.gov/).

Glycosylation and phosphorylation sites were predicted with NetNGlyc 1.0 (https://services.healthtech.dtu.dk/service.php?NetNGlyc-1.0), NetOGlyc 4.0 (http://www.cbs.dtu.dk/services/NetOGlyc/) (51), NetPhos 3.1 (http://www.cbs.dtu.dk/services/NetPhos/) (64), and PhosphoSitePlus (https://www.phosphosite.org) (65) respectively. The dbPTM database (http://dbptm.mbc.nctu.edu.tw/) (66, 67) was queried together with PhosphoSitePlus to retrieve experimentally supported post-translational modifications.

Ser-Gly-X-Gly, X being any amino acid residue except proline (68) and Glu/Asp-x-Ser-Gly (69) sequences, corresponding to glycosaminoglycan (GAG) attachment sites were searched with PATTINPROT (https://npsa-prabi.ibcp.fr) (70). The Ser-Gly pattern was searched manually in SNED1 sequence.

Disulfide bond forming cysteines and ternary cysteine classification were performed using DISULFIND (34) and the DiANNA 1.1: web server (71) respectively.

### Molecular modeling

Two template-free, *ab-initio* protein structure prediction methods, I-TASSER, also integrating a threading approach (36), and QUARK (37) were used to generate models of fl-SNED1 and of the NIDO domain. RaptorX (72), integrating a remote templates strategy, was also used to build a 3D model of fl-SNED1. All the models were refined with MolRefiner (73).

### Predicted interaction network of SNED1

Domain-domain interactions were retrieved using 3did (https://3did.irbbarcelona.org/) (74) to identify putative domains able to interact with SNED1. Queries were performed using the following Pfam (75) domains identifiers: NIDO: PF06119, EGF: PF00008, EGF_CA: PF07645, FOLN: PF09289, Sushi: PF00084, fn3: PF00041. Interactions networks were built with Cytoscape 3.7.2 (76).

MatrixDB (42) and IntAct (43) databases were queried to retrieve experimentally-supported interactions mediated by SNED1. The SNED1 gene identifier was used to query PrePPI (Predicting Protein-Protein Interactions by Combining Structural and non-Structural Information (77)), FpClass, a data mining-based method for proteome-wide protein-protein prediction (78), the Integrated Interactions Database (IID), which provides context-specific physical protein-protein interactions (79), and Struct2Net, a web service predicting protein-protein interactions using a structure-based approach (80). SNED1 sequence (residues 25-1413) was used to query HOMCOS server (http://homcos.pdbj.org/) to predict interacting protein pairs and interacting sites by homology modeling of complex structures (81, 82).

### Reactome Pathway Analysis

The 114 proteins predicted to interact with SNED1 were input as a dataset into the Reactome pathway algorithm (https://reactome.org/) (52, 83). In brief, a statistical test determines whether certain Reactome pathways and biological processes are enriched in the submitted dataset. The test produces a probability score corrected for false discovery rate using the Benjamini-Hochberg method.

## ACKNOWLEDGMENTS

SEC-SAXS, SEC-MALLS, and DLS experiments were performed on the SWING beamline (French National Synchrotron Facility SOLEIL, Saint-Aubin, France, experiment n°20181074). The authors are grateful to Aurélien Thureau from SWING beamline and Frank Gondelaud (ICBMS, UMR 5246, University Lyon 1) for their skillful assistance in SEC-SAXS/MALLS experiments and data interpretation. The authors are also grateful for the UMS 3444/US8 Facility (Lyon, France) for providing access to the Chirascan system. AN would also like to thank Dr. Richard Hynes (Massachusetts Institute of Technology) for his kind gift of the anti-SNED1 antibody and for supporting the initial phase of the development of this project.

## AUTHOR CONTRIBUTIONS

SDV conducted experiments (protein expression and purification, biophysical characterization, interaction prediction and molecular modeling), contributed to preparing the figures, and wrote the manuscript

MND conducted experiments (protein expression, purification, and biochemical characterization), contributed to preparing the figures, and reviewed the manuscript

AB conducted experiment (cloning of SNED1 constructs, sequence analysis) and experiments, contributed to preparing the figures, and reviewed the manuscript

SRB designed and co-supervised the study, contributed to preparing the figures, and wrote the manuscript

AN conceived, designed, and supervised the study, contributed to preparing the figures, and wrote the manuscript

## ABBREVIATIONS

CD: circular dichroism
DLS: dynamic light scattering
DOC: deoxycholate
ECM: extracellular matrix
fl: full length
EGF: Epidermal growth factor
GAG: glycosaminoglycan
HBS: Hepes buffered saline
MALLS: multi-angle laser light scattering
M_w_: molecular weight
PNGase F: peptide-N-glycosidase F
PTM: post-translational modification
Rg: radius of gyration
Rh: hydrodynamic radius
SEC: size-exclusion chromatography
SDS-PAGE: sodium dodecyl sulfate - polyacrylamide gel electrophoresis
SNED1: Sushi, nidogen and EGF-like domain-containing protein 1
SWING: Small and Wide angle X-ray scattering

## SUPPORTING INFORMATION

**Supplementary Figure S1.** Generation of a rabbit anti-SNED1 antibody

**Supplementary Figure S2.** Prediction of the secondary structure of the N-terminal fragment of SNED1

**Supplementary Figure S3.** Size-exclusion chromatography analysis

**Supplementary Figure S4.** *Ab-initio* 3D models of human fl-SNED1

**Supplementary Figure S5.** Recombinant human SNED1 is not modified by GAG chains

**Supplementary Figure S6.** Computational analysis of the predicted SNED1 interactome

**Supplementary Table S1**. List of disulfide bonds predicted in human SNED1 with DISULFIND (S1A) and DiANNA (S1B).

**Supplementary Table S2**. List of predicted (S2A) and experimentally supported (S2B) glycosylation and phosphorylation sites of human SNED1.

**Supplementary Table S3.** List of domain-domain interactions retrieved from 3did using the SNED1 domains as queries.

**Supplementary Table S4**. Lists of predicted partners of human fl-SNED1 using different algorithms.

**Supplementary Table S5**. List of primers used to clone SNED1 constructs.

**Supplementary File S1**: pdb file of the coordinates of the best model of human NIDO generated by QUARK.

**Supplementary File S2**: pdb file of the coordinates of the best model of human fl-SNED1 generated by I-TASSER.

**Supplementary File S3**: pdb file of the coordinates of the best model of human fl-SNED1 generated by RaptorX.

## Footnotes

1> *Unpublished data from the Naba lab*

## REFERENCES

1. Hynes, R. O., and Yamada, K. M. (2012) Extracellular Matrix Biology., Cold Spring Harbor Perspectives in Biology, Cold Spring Harbor Laboratory Press, Cold Spring Harbor, NY

2. Cruz Walma, D. A., and Yamada, K. M. (2020) The extracellular matrix in development. Development. 10.1242/dev.175596

3. Labat-Robert, J., Robert, A.-M., and Robert, L. (2012) Aging of the extracellular matrix. Médecine & Longévité. 4, 3–32

4. Dzamba, B. J., and DeSimone, D. W. (2018) Extracellular Matrix (ECM) and the Sculpting of Embryonic Tissues. Curr. Top. Dev. Biol. 130, 245–274

5. Bonnans, C., Chou, J., and Werb, Z. (2014) Remodelling the extracellular matrix in development and disease. Nature Reviews Molecular Cell Biology. 15, 786–801

6. Karamanos, N. K., Theocharis, A. D., Neill, T., and Iozzo, R. V. (2019) Matrix modeling and remodeling: A biological interplay regulating tissue homeostasis and diseases. Matrix Biology. 75-76, 1–11

7. Theocharis, A. D., Manou, D., and Karamanos, N. K. (2019) The extracellular matrix as a multitasking player in disease. The FEBS Journal. 286, 2830–2869

8. Bruckner, P. (2009) Suprastructures of extracellular matrices: paradigms of functions controlled by aggregates rather than molecules. Cell Tissue Res. 339, 7

9. Hynes, R. O. (2009) The extracellular matrix: not just pretty fibrils. Science. 326, 1216–9

10. Adams, J. C., and Engel, J. (2007) Bioinformatic analysis of adhesion proteins. Methods Mol. Biol. 370, 147–172

11. Hohenester, E., and Engel, J. (2002) Domain structure and organisation in extracellular matrix proteins. Matrix Biology. 21, 115–128

12. Whittaker, C. A., Bergeron, K. F., Whittle, J., Brandhorst, B. P., Burke, R. D., and Hynes, R. O. (2006) The echinoderm adhesome. Developmental biology. 300, 252–66

13. Naba, A., Clauser, K. R., Hoersch, S., Liu, H., Carr, S. A., and Hynes, R. O. (2012) The matrisome: in silico definition and in vivo characterization by proteomics of normal and tumor extracellular matrices. Mol Cell Proteomics. 11, M111.014647

14. Ricard-Blum, S. (2011) The Collagen Family. Cold Spring Harbor Perspectives in Biology. 3, a004978–a004978

15. Iozzo, R. V., and Schaefer, L. (2015) Proteoglycan form and function: A comprehensive nomenclature of proteoglycans. Matrix Biol. 42, 11–55

16. Hynes, R. O., and Naba, A. (2012) Overview of the Matrisome -An Inventory of Extracellular Matrix Constituents and Functions. Cold Spring Harbor Perspectives in Biology. 4, a004903–a004903

17. Naba, A., Hoersch, S., and Hynes, R. O. (2012) Towards definition of an ECM parts list: an advance on GO categories. Matrix Biol. 31, 371–372

18. Leimeister, C., Schumacher, N., Diez, H., and Gessler, M. (2004) Cloning and expression analysis of the mouse stroma marker Snep encoding a novel nidogen domain protein. Dev. Dyn. 230, 371–377

19. Barque, A., Jan, K., Fuente, E. D. L., Nicholas, C. L., Hynes, R. O., and Naba, A. (2020) Knockout of the gene encoding the extracellular matrix protein Sned1 results in craniofacial malformations and early neonatal lethality. bioRxiv. 10.1101/440081

20. Naba, A., Clauser, K. R., Lamar, J. M., Carr, S. A., and Hynes, R. O. (2014) Extracellular matrix signatures of human mammary carcinoma identify novel metastasis promoters. eLife. 3, e01308

21. Letunic, I., and Bork, P. (2018) 20 years of the SMART protein domain annotation resource. Nucleic Acids Res. 46, D493–D496

22. Özbek, S., Balasubramanian, P. G., Chiquet-Ehrismann, R., Tucker, R. P., and Adams, J. C. (2010) The Evolution of Extracellular Matrix. Mol Biol Cell. 21, 4300–4305

23. Hynes, R. O. (2012) The evolution of metazoan extracellular matrix. J Cell Biol. 196, 671–679

24. Pozzi, A., Yurchenco, P. D., and Iozzo, R. V. (2017) The nature and biology of basement membranes. Matrix Biol. 57-58, 1–11

25. Timpl, R., Fujiwara, S., Dziadek, M., Aumailley, M., Weber, S., and Engel, J. (1984) Laminin, proteoglycan, nidogen and collagen IV: structural models and molecular interactions. Ciba Found. Symp. 108, 25–43

26. Goodyear, R. J., and Richardson, G. P. (2002) Extracellular matrices associated with the apical surfaces of sensory epithelia in the inner ear: molecular and structural diversity. J. Neurobiol. 53, 212–227

27. Legan, P. K., Rau, A., Keen, J. N., and Richardson, G. P. (1997) The mouse tectorins. Modular matrix proteins of the inner ear homologous to components of the sperm-egg adhesion system. J. Biol. Chem. 272, 8791–8801

28. Kim, D.-K., Kim, J. A., Park, J., Niazi, A., Almishaal, A., and Park, S. (2019) The release of surface-anchored α-tectorin, an apical extracellular matrix protein, mediates tectorial membrane organization. Sci Adv. 5, eaay6300

29. Izumi, Y., Yanagihashi, Y., and Furuse, M. (2012) A novel protein complex, Mesh-Ssk, is required for septate junction formation in the Drosophila midgut. J Cell Sci. 125, 4923–4933

30. Senapati, S., Gnanapragassam, V. S., Moniaux, N., Momi, N., and Batra, S. K. (2012) Role of MUC4-NIDO domain in the MUC4-mediated metastasis of pancreatic cancer cells. Oncogene. 31, 3346–3356

31. Izzi, V., Davis, M. N., and Naba, A. (2020) Pan-Cancer Analysis of the Genomic Alterations and Mutations of the Matrisome. Cancers. 12, 2046

32. Wierzbicka-Patynowski, I., Mao, Y., and Schwarzbauer, J. E. (2004) Analysis of Fibronectin Matrix Assembly. Current Protocols in Cell Biology. 25, 10.12.1-10.12.10

33. Taylor-Weiner, H., Schwarzbauer, J. E., and Engler, A. J. (2013) Defined Extracellular Matrix Components are Necessary for Definitive Endoderm Induction. Stem Cells. 10.1002/stem.1453

34. Montgomerie, S., Cruz, J. A., Shrivastava, S., Arndt, D., Berjanskii, M., and Wishart, D. S. (2008) PROTEUS2: a web server for comprehensive protein structure prediction and structure-based annotation. Nucleic Acids Res. 36, W202–W209

35. Ceroni, A., Passerini, A., Vullo, A., and Frasconi, P. (2006) DISULFIND: a disulfide bonding state and cysteine connectivity prediction server. Nucleic Acids Res. 34, W177–W181

36. Yang, J., and Zhang, Y. (2015) I-TASSER server: new development for protein structure and function predictions. Nucleic Acids Res. 43, W174–W181

37. Xu, D., and Zhang, Y. (2013) Toward optimal fragment generations for ab initio protein structure assembly. Proteins. 81, 229–239

38. Reily, C., Stewart, T. J., Renfrow, M. B., and Novak, J. (2019) Glycosylation in health and disease. Nature Reviews Nephrology. 15, 346–366

39. Klement, E., and Medzihradszky, K. F. (2017) Extracellular Protein Phosphorylation, the Neglected Side of the Modification. Mol Cell Proteomics. 16, 1–7

40. Goldberg, M. (2012) Phosphorylated Extracellular Matrix Proteins of Bone and Dentin, Bentham Science Publishers

41. Yalak, G., and Olsen, B. R. (2015) Proteomic database mining opens up avenues utilizing extracellular protein phosphorylation for novel therapeutic applications. Journal of Translational Medicine. 13, 125

42. Clerc, O., Deniaud, M., Vallet, S. D., Naba, A., Rivet, A., Perez, S., Thierry-Mieg, N., and Ricard-Blum, S. (2019) MatrixDB: integration of new data with a focus on glycosaminoglycan interactions. Nucleic Acids Res. 47, D376–D381

43. Orchard, S., Ammari, M., Aranda, B., Breuza, L., Briganti, L., Broackes-Carter, F., Campbell, N. H., Chavali, G., Chen, C., del-Toro, N., Duesbury, M., Dumousseau, M., Galeota, E., Hinz, U., Iannuccelli, M., Jagannathan, S., Jimenez, R., Khadake, J., Lagreid, A., Licata, L., Lovering, R. C., Meldal, B., Melidoni, A. N., Milagros, M., Peluso, D., Perfetto, L., Porras, P., Raghunath, A., Ricard-Blum, S., Roechert, B., Stutz, A., Tognolli, M., van Roey, K., Cesareni, G., and Hermjakob, H. (2014) The MIntAct project--IntAct as a common curation platform for 11 molecular interaction databases. Nucleic Acids Res. 42, D358–363

44. Stellato, C., Nassa, G., Tarallo, R., Giurato, G., Ravo, M., Rizzo, F., Marchese, G., Alexandrova, E., Cordella, A., Baumann, M., Nyman, T. A., Weisz, A., and Ambrosino, C. (2015) Identification of cytoplasmic proteins interacting with unliganded estrogen receptor α and β in human breast cancer cells. Proteomics. 15, 1801–1807

45. Campbell, I. D., and Humphries, M. J. (2011) Integrin Structure, Activation, and Interactions. Cold Spring Harbor Perspectives in Biology. 3, a004994–a004994

46. Midwood, K., and Orend, G. (2015) Special issue of Cell Adhesion & Migration on Tenascins: Defining their role in tissue homeostasis and cancer. Cell Adhesion & Migration. 9, 1–3

47. Takeuchi, H., Yu, H., Hao, H., Takeuchi, M., Ito, A., Li, H., and Haltiwanger, R. S. (2017) O-Glycosylation modulates the stability of epidermal growth factor-like repeats and thereby regulates Notch trafficking. J. Biol. Chem. 292, 15964–15973

48. Sethi, M. K., Buettner, F. F. R., Krylov, V. B., Takeuchi, H., Nifantiev, N. E., Haltiwanger, R. S., Gerardy-Schahn, R., and Bakker, H. (2010) Identification of glycosyltransferase 8 family members as xylosyltransferases acting on O-glucosylated notch epidermal growth factor repeats. J. Biol. Chem. 285, 1582–1586

49. Teng, Y., Liu, Q., Ma, J., Liu, F., Han, Z., Wang, Y., and Wang, W. (2006) Cloning, expression and characterization of a novel human CAP10-like gene hCLP46 from CD34+ stem/progenitor cells. Gene. 371, 7–15

50. Wang, Y., Shao, L., Shi, S., Harris, R. J., Spellman, M. W., Stanley, P., and Haltiwanger, R. S. (2001) Modification of Epidermal Growth Factor-Like Repeats With O-fucose. Molecular Cloning and Expression of a Novel GDP-fucose Protein O-fucosyltransferase. J. Biol. Chem. 276, 40338–40345

51. Steentoft, C., Vakhrushev, S. Y., Joshi, H. J., Kong, Y., Vester-Christensen, M. B., Schjoldager, K. T.-B. G., Lavrsen, K., Dabelsteen, S., Pedersen, N. B., Marcos-Silva, L., Gupta, R., Paul Bennett, E., Mandel, U., Brunak, S., Wandall, H. H., Levery, S. B., and Clausen, H. (2013) Precision mapping of the human O-GalNAc glycoproteome through SimpleCell technology. EMBO J. 32, 1478–1488

52. Jassal, B., Matthews, L., Viteri, G., Gong, C., Lorente, P., Fabregat, A., Sidiropoulos, K., Cook, J., Gillespie, M., Haw, R., Loney, F., May, B., Milacic, M., Rothfels, K., Sevilla, C., Shamovsky, V., Shorser, S., Varusai, T., Weiser, J., Wu, G., Stein, L., Hermjakob, H., and D’Eustachio, P. (2020) The reactome pathway knowledgebase. Nucleic Acids Res. 48, D498–D503

53. Zhu, Y., Zhang, J.-J., Peng, Y.-P., Liu, X., Xie, K.-L., Tang, J., Jiang, K.-R., Gao, W.-T., Tian, L., Zhang, K., Xu, Z.-K., and Miao, Y. (2017) NIDO, AMOP and vWD domains of MUC4 play synergic role in MUC4 mediated signaling. Oncotarget. 8, 10385–10399

54. Weissmueller, S., Manchado, E., Saborowski, M., Morris, J. P., Wagenblast, E., Davis, C. A., Moon, S.-H., Pfister, N. T., Tschaharganeh, D. F., Kitzing, T., Aust, D., Markert, E. K., Wu, J., Grimmond, S. M., Pilarsky, C., Prives, C., Biankin, A. V., and Lowe, S. W. (2014) Mutant p53 drives pancreatic cancer metastasis through cell-autonomous PDGF receptor beta signaling. Cell. 157, 382–394

55. Hamidi, H., and Ivaska, J. (2018) Every step of the way: integrins in cancer progression and metastasis. Nature Reviews Cancer. 10.1038/s41568-018-0038-z

56. Villafuerte, B. C., and Kaytor, E. N. (2005) An insulin-response element-binding protein that ameliorates hyperglycemia in diabetes. J. Biol. Chem. 280, 20010–20020

57. Villafuerte, B. C., Phillips, L. S., Rane, M. J., and Zhao, W. (2004) Insulin-response Element-binding Protein 1. A novel Akt substrate involved in transcriptional action of insulin. J. Biol. Chem. 279, 36650–36659

58. Vallet, S. D., Miele, A. E., Uciechowska-Kaczmarzyk, U., Liwo, A., Duclos, B., Samsonov, S. A., and Ricard-Blum, S. (2018) Insights into the structure and dynamics of lysyl oxidase propeptide, a flexible protein with numerous partners. Sci Rep. 8, 11768

59. Gondelaud, F., Connes, P., Nader, E., Renoux, C., Fort, R., Gauthier, A., Joly, P., and Ricard-Blum, S. (2020) Sialic acids rather than glycosaminoglycans affect normal and sickle red blood cell rheology by binding to four major sites on fibrinogen. Am. J. Hematol. 10.1002/ajh.25718

60. Whitmore, L., and Wallace, B. A. (2008) Protein secondary structure analyses from circular dichroism spectroscopy: Methods and reference databases. Biopolymers. 89, 392–400

61. Compton, L. A., and Johnson, W. C. (1986) Analysis of protein circular dichroism spectra for secondary structure using a simple matrix multiplication. Anal. Biochem. 155, 155–167

62. Nygaard, M., Kragelund, B. B., Papaleo, E., and Lindorff-Larsen, K. (2017) An Efficient Method for Estimating the Hydrodynamic Radius of Disordered Protein Conformations. Biophys. J. 113, 550–557

63. Liu, W., Xie, Y., Ma, J., Luo, X., Nie, P., Zuo, Z., Lahrmann, U., Zhao, Q., Zheng, Y., Zhao, Y., Xue, Y., and Ren, J. (2015) IBS: an illustrator for the presentation and visualization of biological sequences. Bioinformatics. 31, 3359–3361

64. Blom, N., Gammeltoft, S., and Brunak, S. (1999) Sequence and structure-based prediction of eukaryotic protein phosphorylation sites. J. Mol. Biol. 294, 1351–1362

65. Hornbeck, P. V., Zhang, B., Murray, B., Kornhauser, J. M., Latham, V., and Skrzypek, E. (2015) PhosphoSitePlus, 2014: mutations, PTMs and recalibrations. Nucleic Acids Res. 43, D512–520

66. Huang, K.-Y., Su, M.-G., Kao, H.-J., Hsieh, Y.-C., Jhong, J.-H., Cheng, K.-H., Huang, H.-D., and Lee, T.-Y. (2016) dbPTM 2016: 10-year anniversary of a resource for post-translational modification of proteins. Nucleic Acids Res. 44, D435–446

67. Huang, K.-Y., Lee, T.-Y., Kao, H.-J., Ma, C.-T., Lee, C.-C., Lin, T.-H., Chang, W.-C., and Huang, H.-D. (2019) dbPTM in 2019: exploring disease association and cross-talk of post-translational modifications. Nucleic Acids Res. 47, D298–D308

68. Yu, Y., and Linhardt, R. J. (2018) Xylosyltransferase 1 and the GAG Attachment Site. Structure. 26, 797–799

69. Okahata, S., Yamamoto, R., Yamakoshi, Y., and Fukae, M. (2011) A Large Chondroitin Sulfate Proteoglycan, Versican, in Porcine Predentin. J Oral Biosci. 53, 72–81

70. Combet, C., Blanchet, C., Geourjon, C., and Deléage, G. (2000) NPS@: network protein sequence analysis. Trends Biochem. Sci. 25, 147–150

71. Ferrè, F., and Clote, P. (2006) BTW: a web server for Boltzmann time warping of gene expression time series. Nucleic Acids Res. 34, W482–485

72. Källberg, M., Wang, H., Wang, S., Peng, J., Wang, Z., Lu, H., and Xu, J. (2012) Template-based protein structure modeling using the RaptorX web server. Nat Protoc. 7, 1511–1522

73. Xu, D., and Zhang, Y. (2011) Improving the physical realism and structural accuracy of protein models by a two-step atomic-level energy minimization. Biophys. J. 101, 2525–2534

74. Mosca, R., Céol, A., Stein, A., Olivella, R., and Aloy, P. (2014) 3did: a catalog of domain-based interactions of known three-dimensional structure. Nucleic Acids Res. 42, D374–D379

75. Finn, R. D., Miller, B. L., Clements, J., and Bateman, A. (2014) iPfam: a database of protein family and domain interactions found in the Protein Data Bank. Nucleic Acids Res. 42, D364–373

76. Shannon, P., Markiel, A., Ozier, O., Baliga, N. S., Wang, J. T., Ramage, D., Amin, N., Schwikowski, B., and Ideker, T. (2003) Cytoscape: a software environment for integrated models of biomolecular interaction networks. Genome Res. 13, 2498–2504

77. Zhang, Q. C., Petrey, D., Garzón, J. I., Deng, L., and Honig, B. (2013) PrePPI: a structure-informed database of protein-protein interactions. Nucleic Acids Res. 41, D828–D833

78. Kotlyar, M., Pastrello, C., Pivetta, F., Lo Sardo, A., Cumbaa, C., Li, H., Naranian, T., Niu, Y., Ding, Z., Vafaee, F., Broackes-Carter, F., Petschnigg, J., Mills, G. B., Jurisicova, A., Stagljar, I., Maestro, R., and Jurisica, I. (2015) In silico prediction of physical protein interactions and characterization of interactome orphans. Nat. Methods. 12, 79–84

79. Kotlyar, M., Pastrello, C., Malik, Z., and Jurisica, I. (2019) IID 2018 update: context-specific physical protein-protein interactions in human, model organisms and domesticated species. Nucleic Acids Res. 47, D581–D589

80. Singh, R., Park, D., Xu, J., Hosur, R., and Berger, B. (2010) Struct2Net: a web service to predict protein-protein interactions using a structure-based approach. Nucleic Acids Res. 38, W508–515

81. Fukuhara, N., and Kawabata, T. (2008) HOMCOS: a server to predict interacting protein pairs and interacting sites by homology modeling of complex structures. Nucleic Acids Res. 36, W185–189

82. Kawabata, T. (2016) HOMCOS: an updated server to search and model complex 3D structures. J Struct Funct Genomics. 17, 83–99

83. Fabregat, A., Sidiropoulos, K., Viteri, G., Marin-Garcia, P., Ping, P., Stein, L., D’Eustachio, P., and Hermjakob, H. (2018) Reactome diagram viewer: data structures and strategies to boost performance. Bioinformatics. 34, 1208–1214

84. Schramm, A., Bignon, C., Brocca, S., Grandori, R., Santambrogio, C., and Longhi, S. (2019) An arsenal of methods for the experimental characterization of intrinsically disordered proteins - How to choose and combine them? Arch. Biochem. Biophys. 10.1016/j.abb.2019.07.020

