## Supplementary Figures for "Computational and experimental characterization of the novel ECM glycoprotein SNED1 and prediction of its interactome"

**Supplementary Figure S1. Generation of a rabbit anti-SNED1 antibody**

**A.**

```
hSNED1    29ADFYPFGAERGDA41
mSned1    29ADFYPFGTKRGDT41
            *******__***_
```

**B.**

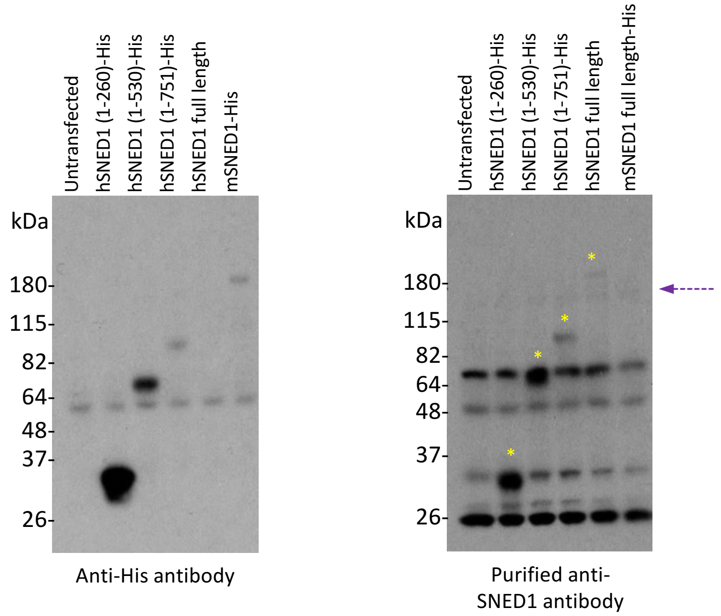


**A**. Sequence of the peptide from human SNED1 (hSNED1) used for immunization, and alignment with the murine sequence (mSned1).

**B**. 293T cells were transiently transfected with different cDNA constructs encoding His-tagged fragments of human SNED1 (corresponding amino acid residues indicated in parentheses), full-length untagged human SNED1, or full-length His-tagged mouse Sned1 (*see Materials and Methods*). Cells were lysed in 3X Laemmli buffer containing 100 mM DTT and proteins resolved by SDS-PAGE. Western blots were performed using a mouse anti-His antibody (Invitrogen #46-0693 used at 1 µg/mL; left panel) or the purified rabbit polyclonal anti-SNED1 antibody we generated (rabbit 7580, antibody eluted with a 0.2 M glycine solution pH 3 and used at 0.5 µg/ml; right panel) as primary antibodies and HRP-coupled anti-mouse or anti-rabbit secondary antibodies respectively.

The rabbit polyclonal anti-SNED1 antibody we generated recognizes specifically human fl-SNED1 and its fragments (yellow stars) but not murine Sned1. Note that the band present in all 6 lanes between the 115 kDa and 180 kDa markers (purple arrow) and detected with the anti-SNED1 antibody, but not the anti-His antibody, may correspond to the endogenous SNED1 expressed by 293T cells.

**Supplementary Figure S2. Prediction of the secondary structure of the N-terminal fragment of SNED1**

```
 25  AVALADFYPF GAERGDAVTP KQDDGGSGLR PLSVPFPFFG AEHSGLYVNN
 75  NGIISFLKEV SQFTPVAFPI AKDRCVVAAF WADVDNRRAG DVYYREATDP
125  AMLRRATEDV RHYFPELLDF NATWVFVATW YRVTFFGGSS SSPVNTFQTV
175  LITDGKLSFT IFNYESIVWT TGTHASSGGN ATGLGGIAAQ AGFNAGDGQR
225  YFSIPGSRTA DMAEVETTTN VGVPGRWAFR IDDAQV
```

| |
|---|
| 9.3% helix<br>33% β-strand |

Prediction of the secondary structures found in the N-terminal fragment of human SNED1 using Proteus2.

Number of residues read in: 236

No homolog was found

Number of sequence alignments used for ab-initio predictions: 49

Overall confidence value: 78.**9**%

Predicted % Helix content: 9 % (22 residues)

Predicted % Beta sheet content: 33 % (**78 residues**)

Predicted % Coil content: 58 % (**136 residues**)

### Supplementary Figure S3. Size-exclusion chromatography analysis

**A.**

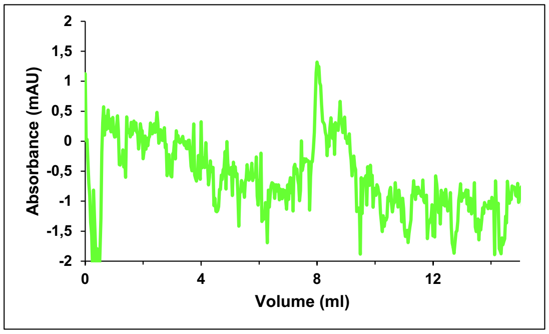


**B.**

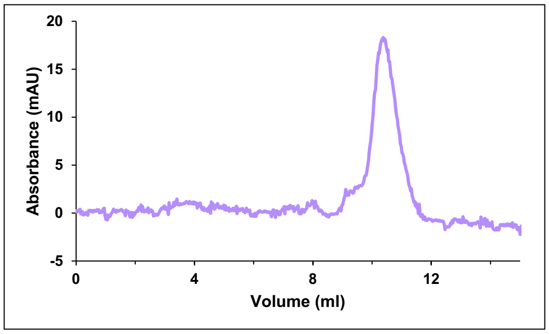


**A-B.** Size-Exclusion Chromatography analysis of fl-SNED1 (**A**) and the N-terminal fragment of SNED1 (**B**). 100 µl of each protein (N-terminal fragment at 11.7 µM; fl-SNED1 at 1.3 µM), diluted in HBS were injected with an Äkta purifier system (GE Healthcare) over a Superdex 75 Increase 10-300 GL (GE Healthcare) and a Superdex 200 Increase 10/300 GL (GE Healthcare) respectively at a flow rate of 0.4 ml/min at 4°C. Calibrations were performed according to manufacturer's instructions with the gel filtration calibration kits Low and High Molecular Weight (GE Healthcare GE28-4038-41 and GE28-4038-42).

### Supplementary Figure S4. *Ab-initio* 3D models of human fl-SNED1

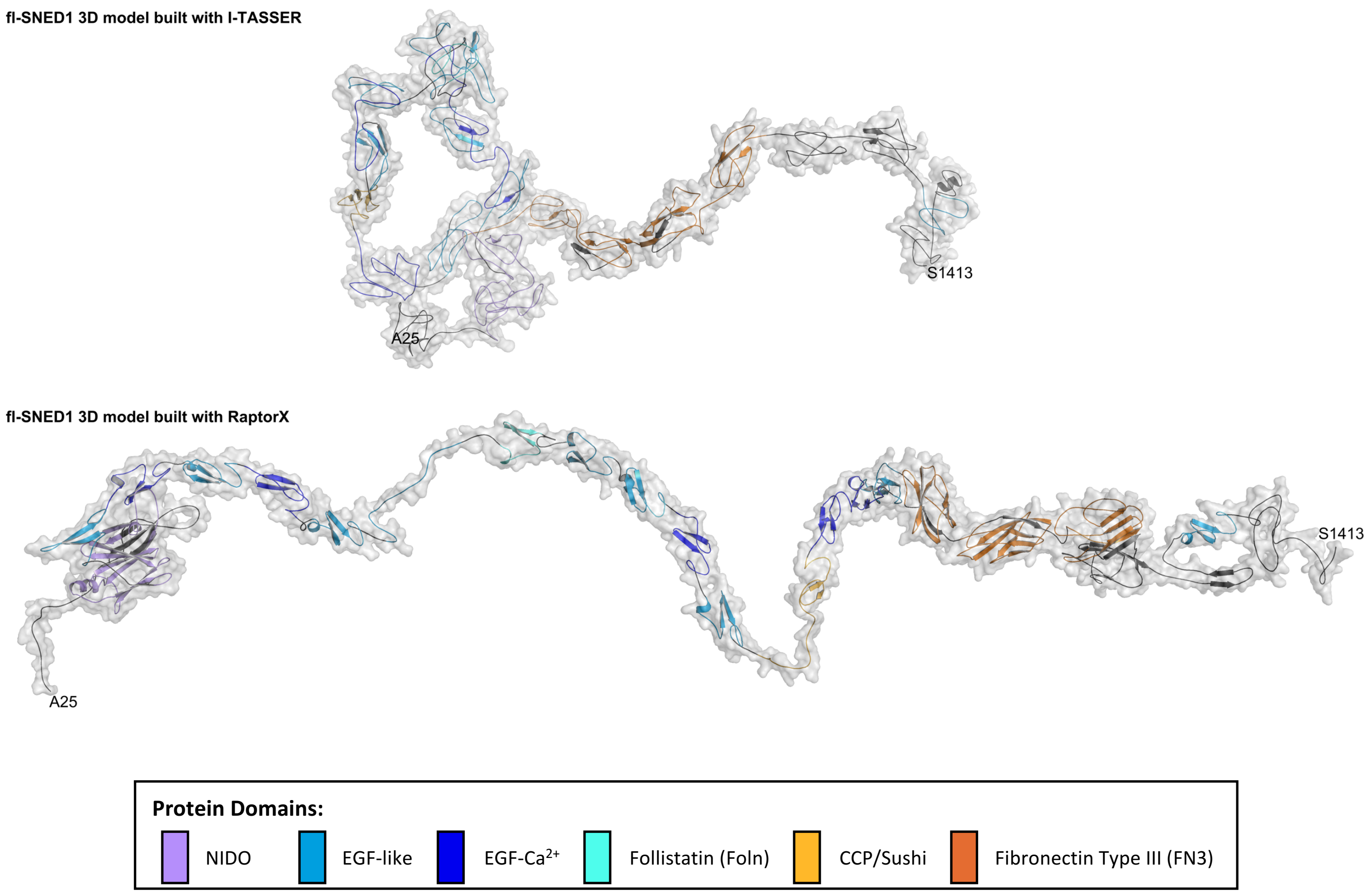


*Ab initio* tentative 3D models of human fl-SNED1 generated by I-TASSER (C-score: - 0.98, TM-score: 0.59, QMEAN -13.79; Upper panel), and by RaptorX (global distance test GDT: 43, un-normalized GDT: 597, QMEAN -4.87; Lower panel) and refined with ModRefiner.

**Supplementary Figure S5. Recombinant human SNED1 is not modified by GAG chains**

**A.**

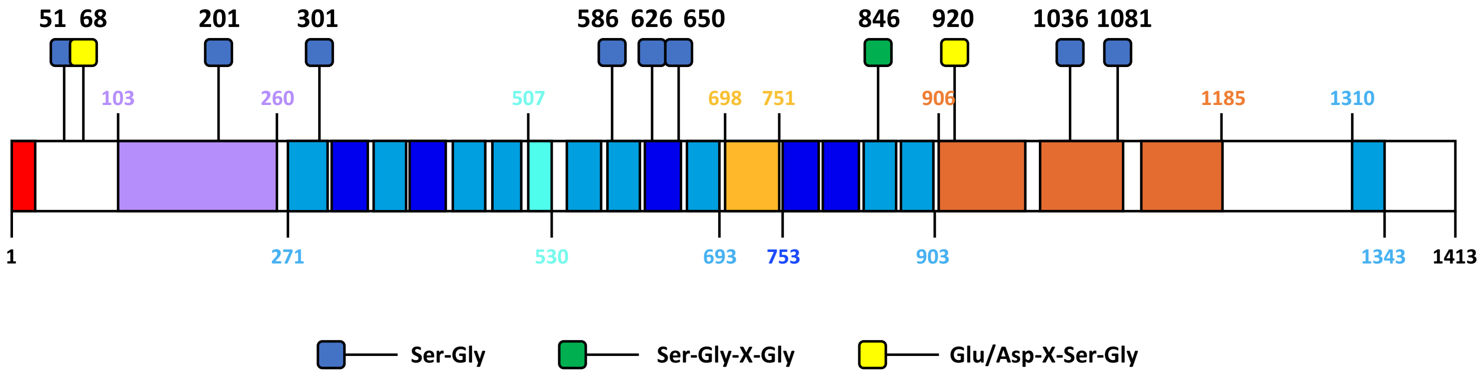


**B.**

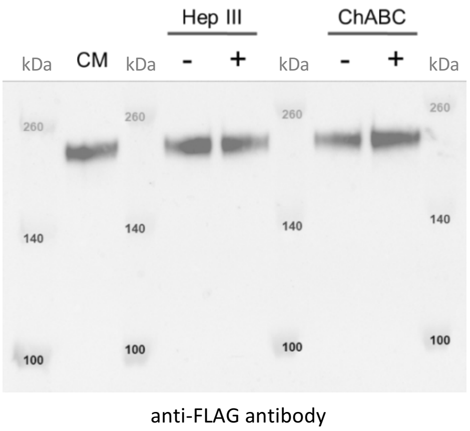


**A.** Predicted sites of glycosaminoglycan attachment in SNED1. Numbers indicate amino acid residue.
**B.** Western blot analysis of the conditioned medium (CM) of 293T cells overexpressing fl-SNED1-FLAG incubated in presence or absence of heparinase III (HepIII), and chondroitinase ABC (ChABC) as described in the Material and Methods section.

**Supplementary Figure S6. Computational analysis of the predicted SNED1 interactome**

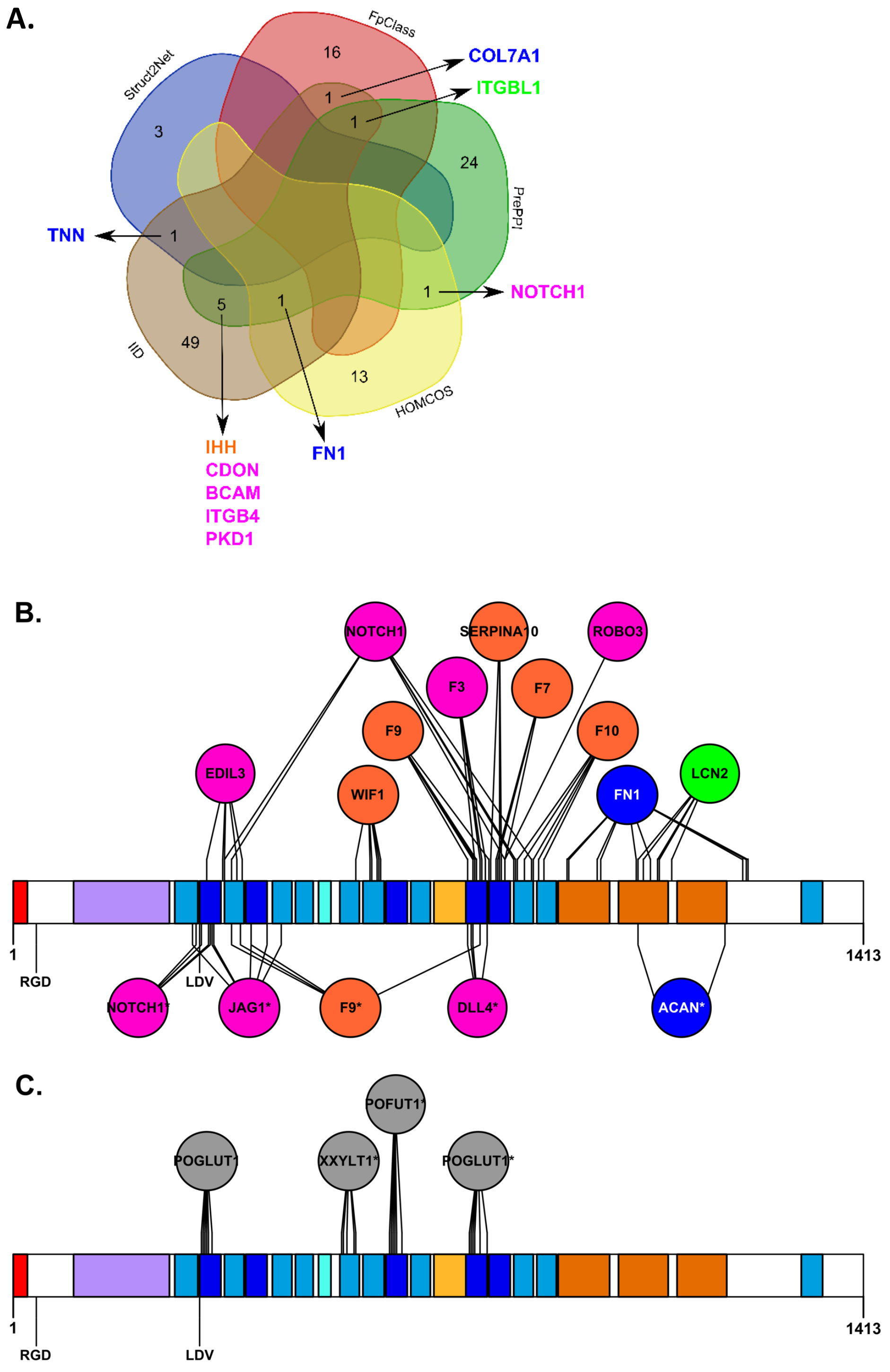


**A**. Venn diagram showing the number of SNED1 partners predicted by various approaches described in the Material and Methods section, and those specifically predicted by a single approach. Proteins predicted to bind to SNED1 by at least two different methods are highlighted. The diagram was drawn with a tool accessible at the following link: http://bioinformatics.psb.ugent.be/webtools/Venn/.

**B-C**. Binding sites of predicted extracellular and transmembrane (**B**) or intracellular (**C**) partners of human SNED1 identified with HOMCOS. Proteins are colored according to the subcellular location of SNED1 partners: blue: core matrisome; orange: matrisome-associated; green: secreted proteins; pink: membrane; grey: intracellular. Note that the gene names corresponding to the human gene names used to build Supplementary Figure S5B and S5C were either directly identified by HOMCOS, or inferred to the human homologs (marked with an asterisk, *). The species of the predicted partners (if different from human) are mentioned in **Supplementary Table S5E**.
