## Supplementary material for "Computational and experimental characterization of the novel ECM glycoprotein SNED1 and prediction of its interactome": Supplementary Table S5_Primers.docx

**Supplementary Table S5: List of primers used in this study**

**Supplementary Table S5A:** Primers used to subclone SNED1 constructs into pMSCV-IRES-Hygromycin.

| **Constructs** | **Primer sequences** |
| --- | --- |
| NIDO-FLAG | Forward primer:  5' TCGAAGATCTGCCACCATGCGGCACGGCGTC 3' |
|  | Reverse primer:  5' CCGTTAACTTA**CTTGTCGTCATCGTCTTTGTAGTC**CACCTGGGCATCATCGATTC 3' |
| fl-SNED1-FLAG | Forward primer:  5' TCGAAGATCTGCCACCATGCGGCACGGCGTC 3' |
|  | Reverse primer:  5' CCGTTAACTTA**CTTGTCGTCATCGTCTTTGTAGTC**AGATTTCTCCAGTGTCTGACTCT 3' |

BglII and HpaI restriction sites are indicated in blue.

ATG (start codon) and STOP codon are indicated in purple.

Sequence of the FLAG tag is indicated in **bold**.

**Supplementary Table S5B:** Primers used to subclone SNED1 constructs into pcDNA5/FRT in frame with a C-terminal 6x-His tag inserted in the vector backbone.

| **Constructs** | **Primer sequences** |
| --- | --- |
| hSNED1 (1-260)-His | Forward primer:  5' ATGCGGCCGGCCCATGCGGCACGGCGTC 3' |
|  | Reverse primer:  5' AGGCGCGCCTCACCTGGGCATCATCGATTCT 3' |
| hSNED1 (1-530)-His | Forward primer:  5' ATGCGGCCGGCCCATGCGGCACGGCGTC 3' |
|  | Reverse primer:  5' AGGCGCGCCTGCACTGGGGAGGCTCACTCCA 3' |
| hSNED1 (1-751)-His | Forward primer:  5' ATGCGGCCGGCCCATGCGGCACGGCGTC 3' |
|  | Reverse primer:  5' AGGCGCGCCTGACGCAGAGGTAGCTCCC 3' |
| mSned1 full length-His | Forward primer:  5' ATGCGGCCGGCCCATGCGCCTCGGCG 3' |
|  | Reverse primer:  5' AGGCGCGCCTCTTTGTTTCCTGTTTGGGTTT 3' |

FseI and AscI restriction sites are indicated in blue.

ATG (start codon) is indicated in purple.
